# Single-allele chromatin folding is organized by recurrent motifs that are reweighted across cell types and states

**DOI:** 10.1101/2024.09.18.613689

**Authors:** Olivier Messina, Jean-Bernard Fiche, Gautham Ganesh, Christel Elkhoury Youhanna, Yasmine Kemkem, Alexandre Mesnager, Jorge Ferrer, Adrian Villalba, Raphael Scharfmann, Ildem Akerman, David J. Hodson, Marie Schaeffer, Marcelo Nollmann

## Abstract

Chromatin folding exhibits extensive cell-to-cell variability, yet whether this variability is dominated by continuously varying conformations or by recurrent organizational patterns remains unclear. Here, we combined high-resolution chromatin tracing with a novel machine learning algorithm to show that chromatin organization can be compactly represented by a limited repertoire of recurrent folding motifs. Using polymer simulations and cohesin depletion experiments, we further show that these motifs arise primarily from loop extrusion. Analysis of distinct tissues reveals that a shared motif repertoire is sufficient to describe diverse structural ensembles, with biological differences arising primarily through changes in motif occupancy and combinatorial usage. More generally, rather than generating entirely new conformations, transcriptional and disease-associated changes, including those occurring during disease onset, act primarily by redistributing motif occupancy within this common repertoire. In particular, gene activation is associated with a shift toward decompacted states and reduced contact frequencies, revealing a quantitative link between transcription and chromatin organization at the single-allele level. Together, these findings reveal that single-allele chromatin variability is structured by a constrained repertoire of recurrent folding motifs whose usage is modulated by regulatory context, providing a new framework to interpret how chromatin organization relates to gene activity.

## Introduction

Eukaryotic chromosomes are organized at multiple scales, with topologically associating domains (TADs)^1–3^ encapsulating genes and most of their *cis*-regulatory elements^4,5^. In mammals, TADs are formed by the combined action of cohesin-mediated loop extrusion^6–8^ and interactions between *cis*-regulatory elements (CRE), such as enhancers (E) and promoters (P)^9–12^, which together shape transcriptional regulation^13^. While these mechanisms have been extensively characterized at the population level, how they sculpt 3D chromatin architecture in single cells, and how this architecture relates to cell-type-specific gene regulation^14^ remains poorly understood.

Single-cell and imaging studies have revealed that TAD architecture is highly heterogeneous^15–22^. Loop extrusion is intrinsically dynamic^7,23^, and CCCTC factor (CTCF)-mediated loops are infrequent and short-lived^24,25^. Enhancer–promoter proximity is also transient and occurs at low frequencies^26–32^. Thus, the principal mechanisms organizing TADs generate a spectrum of structurally diverse conformations across single alleles, consistent with chromatin exploring a high-dimensional conformational landscape. A central question is therefore whether this variability contains reproducible organizational patterns that can be compactly represented despite this underlying conformational heterogeneity. Previous approaches were unable to recover interpretable patterns at regulatory scales^33,34^, because of intrinsic limitations of the method or due to the genomic scales at which chromatin structure was probed. As a result, it remains unclear whether recurrent structural motifs can explain a substantial fraction of the observed variability, and how such organization relates to transcriptional and regulatory processes.

Here, we combine high-resolution chromatin tracing with an unsupervised topic-modeling framework to demonstrate that TAD architecture is built from a constrained ensemble of recurrent folding motifs. Across multiple loci and species, we show that single-allele chromatin conformations can be quantitatively decomposed into combinations of a limited repertoire of motifs. By extending this analysis to multiple tissues, we further show that a shared motif dictionary is sufficient to describe tissue-specific chromatin architectures, with biological differences arising primarily through changes in motif occupancy and combinatorial usage. Next, we integrated polymer simulations and cohesin depletion experiments to show that motifs arise primarily from the action of loop extrusion. Finally, we show that transcription-dependent and disease-associated changes in chromatin organization are largely explained by quantitative redistributions within this common motif repertoire, rather than by the emergence of entirely new structural states, revealing a shared structural grammar underlying TAD organization across biological contexts.

## Results

### Single-allele chromatin conformations can be decomposed into recurrent folding motifs

To investigate whether heterogeneous chromatin conformations contain recurrent organizational patterns that can be compactly represented, we imaged 3D chromatin organization in single alleles within adult mouse pancreas cryosections using high-resolution Hi-M imaging (**Fig. 1a**, Methods). Hi-M is a sequential DNA-FISH imaging technology that enables high-resolution tracing of 3D chromatin architecture across many single alleles in intact tissues^16,27,35^. We focused on a regulatory domain containing the duodenal homeobox 1 gene (*Pdx1*) and its cis-regulatory elements (CREs), a well-characterized locus with defined regulatory inputs in beta cells^36^ (**Fig. 1b**). This system provides a biologically relevant context to resolve how single-allele chromatin variability relates to regulatory state.

**Figure 1.**
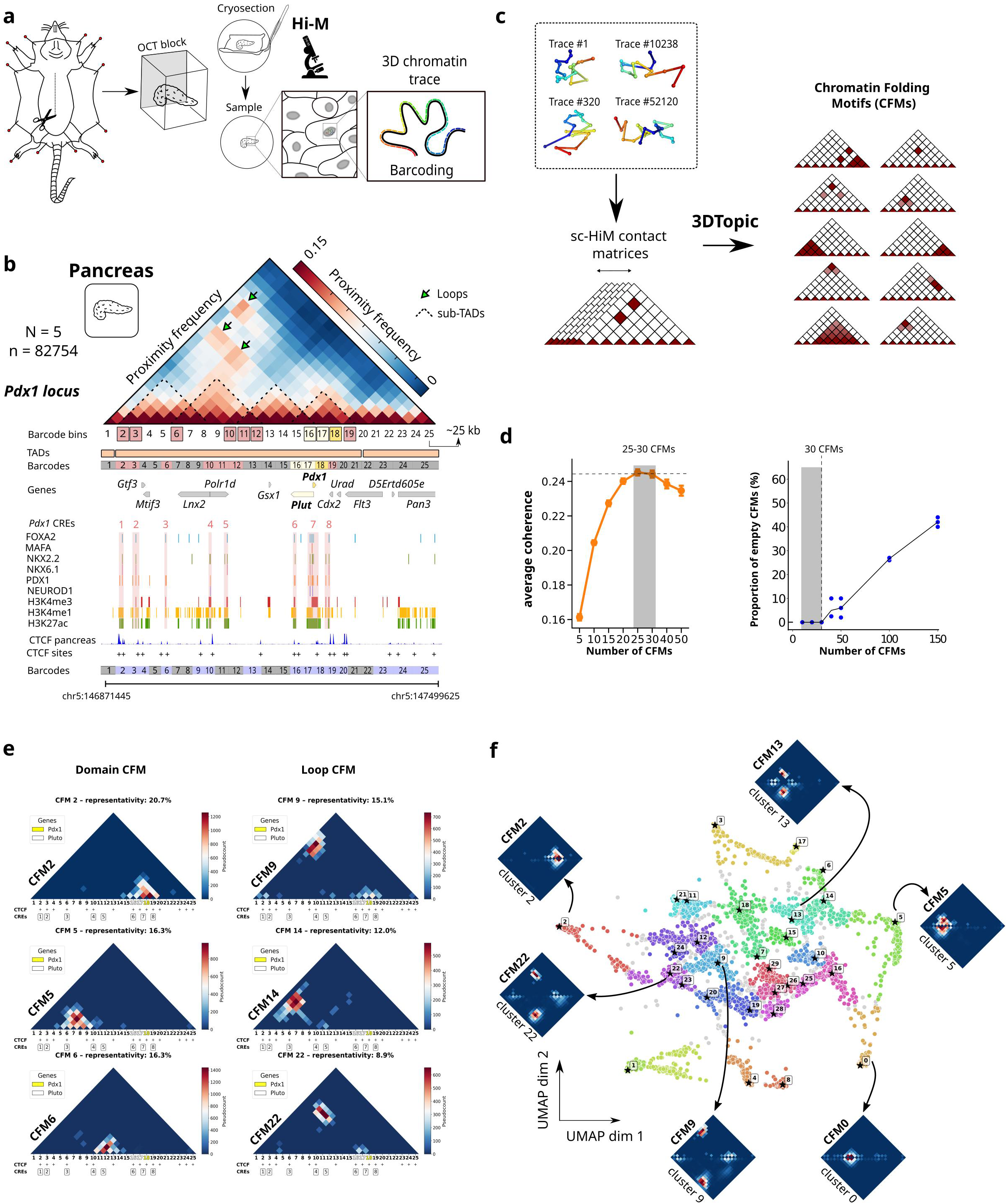
Single-allele chromatin conformations are organized by recurrent folding motifs. **a** Schematic illustrating the imaging-based strategy used to trace chromatin architecture at single-cell level in cryosectioned mouse tissues (Hi-M). **b** Top: Hi-M proximity frequency matrix of the *Pdx1* locus (chr5:146871445-147499625, mm10) from mouse pancreas generated with a distance cutoff of 100 nm, constructed from *n* = 82754 traces (*N* = 5 experiments, 2 different mice). Loops are indicated with arrows, while sub-TADs are marked with dashed lines. Middle: TADs are represented with orange rectangles. Barcodes used for Hi-M sequential imaging are represented as color-coded boxes (pink: presence of CREs, white: *Plut* gene, yellow: *Pdx1* gene). Gene locations are displayed along the locus. Identified CREs are indicated by red numbers. CREs were defined based on the binding of at least two pancreatic TFs. Binding sites of for six pancreatic TFs (FOXA2, MAFA, NKX2.2, NKX6.1, PDX1, and NEUROD1) derived from ChIP-seq in pancreas data are shown below, together with characteristic histone marks associated with active enhancers (H3K4me3, H3K4me1, and H3K27ac). The CTCF profile, derived from CUT&Tag experiments in pancreas, is also plotted, with identified CTCF-binding sites indicated by the “+” signs. Hi-M barcodes are further color-coded according to the presence or absence of CTCF-binding sites (blue: presence, gray: absence). **c** Schematic illustrating 3DTopic workflow and application for single cell Hi-M data. Single cell 3D structures are used to determine a set of Chromatin Folding Motifs (CFMs) and each cell is iteratively assigned to a probability distribution for each CFM (see *Methods*). **d** Left: Distribution of the coherence score (NPMI) for ten separate decompositions with varying numbers of CFMs. The maximum of the coherence score was reached between 25 and 30 CFMs, as highlighted by a gray box (see *Methods*). Right: Proportion of empty CFMs relative to the total number of CFMs. **e** Subsets of CFMs, selected from the complete gallery of 30 CFMs of one decomposition of the Pdx1 locus on pancreatic cells (see *Methods*). CFMs are categorized into two distinct categories according to the structure they describe (Domain and Loop) (see **Fig. S2d**, complete list). **f** UMAP embeddings of CFMs from 200 independent trainings performed with 30 CFMs. CFMs were clustered using the HDBSCAN algorithm, and a single reference CFM (indicated by a star) was identified within each cluster. Surrounding the embedding are example CFMs from the reference decomposition (marked by a star) (top), shown alongside the cluster-average CFM matrix computed from all CFMs within the respective cluster (bottom).

PDX1 is a master regulator of pancreatic development, and in the adult pancreas it is expressed specifically in insulin-producing beta cells, where it is down-regulated in obese and diabetic islets^37^. In mouse and humans, *Pdx1*/*PDX1* transcription is regulated by conserved enhancers bound by key beta-cell specific transcription factors, such as MAFA, FOXA2, NKX6.1, NKX2.2, NEUROD1^36–40^ (**Fig. 1b**). In humans, these enhancers are co-accessible and contact the *PDX1* promoter specifically in beta cells^41^, overlap type 2 diabetes (T2D)-associated variants^36,42^, and exhibit altered chromatin accessibility in T2D islets^41^. PDX1 also participates in an autoregulatory loop that reinforces its expression in mature beta cells^43^, and its transcription can be affected by the beta cell-specific long non-coding RNA Plut which also changes local chromatin structure around *PDX1* and its proximal enhancers^44^. Thus, mapping single-allele 3D chromatin organization at the *Pdx1* locus provides a powerful, disease relevant model in which chromatin organization, transcriptional activity, and disease-associated perturbations can be examined at single-allele resolution.

To characterize how chromatin conformations are distributed across single alleles at this locus, we generated Hi-M chromatin tracing datasets covering a 625-kb region surrounding the *Pdx1* gene using 25 primary barcodes, each spanning on average 25 kb (**Fig. 1b**, Supplementary Data 1). This design represented a tradeoff between genomic resolution and genomic coverage, providing sufficient resolution to image the entire TAD containing *Pdx1* while resolving interactions between *Pdx1* and its distal CREs. Image acquisition, analysis and validation were performed as previously described^27,45^ (**Figs. S1a-d**, and Methods). Different replicates were highly reproducible, as indicated by Pearson’s correlation analysis (**Figs. S1e-f**). The proximity frequency and ensemble pairwise distance (PWD) maps of the *Pdx1* locus display a single TAD with multiple overlapping loops (**Figs. 1b, S1g**). Notably, these structural features are also clearly distinguishable in ensemble Hi-C maps for the same locus (**Fig. S1h**), which correlate well with Hi-M proximity frequencies (**Fig. S1i**). While these ensemble measurements capture the average organization of the locus, individual chromatin traces display substantial variability in pairwise contacts across single alleles (**Fig. 1c**), reflecting the heterogeneous conformational landscape explored by this regulatory domain.

To determine whether this heterogeneity contains recurrent organizational patterns that can be compactly represented despite the underlying conformational variability, we developed 3DTopic, an unsupervised machine learning approach (see *Methods*)^46^. Motivated by the need for an approach robust to missing values, common in high-resolution DNA tracing data, we used latent Dirichlet allocation (LDA), a probabilistic model originally developed for natural language processing that represents each document as a mixture of hidden “topics” and each topic as a characteristic distribution of words, thereby uncovering the latent structure of a corpus of documents (**Fig. 1c**). Similarly, 3DTopic uses the LDA formalism and treats each single-allele contact map as a ‘document’, and individual contacts as ‘words’, and is natively robust to missing values^46^. Topic modeling was used in the past to analyze single-cell Hi-C data^47^, however the low genomic resolution of this method (∼500 kb) limited their applicability for resolving regulatory-scale chromatin organization within TADs.

Notably, by applying 3DTopic to *Pdx1* Hi-M dataset, we find that chromatin conformations can be decomposed into a limited set of recurrent patterns of contacts that co-occur across single alleles, which we refer to as chromatin folding motifs (**CFMs**) (**Fig. 1c**). In this representation, each chromatin trace is described as a mixture of CFMs with associated weights, allowing the heterogeneous conformational ensemble to be decomposed into a compact set of recurrent structural patterns together with their per-allele contributions.

3DTopic operates on single-allele proximity maps, obtained by binarizing pairwise distance maps using a fixed distance cutoff (*Methods*). This step defines the physical interaction scale represented in the input maps rather than a free parameter of the topic model itself. We chose a cut-off distance of 100 nm because it maximizes the correlation between Hi-M proximity frequencies and ensemble Hi-C contact maps at the *Pdx1* locus (**Fig. S1i**), thereby matching the Hi-M contacts used for decomposition with the physical interaction scale most comparable to Hi-C.

To evaluate the robustness of 3DTopic and determine appropriate model parameters, we performed a series of benchmarking analyses assessing reproducibility, reconstruction accuracy, and sensitivity to model complexity. The optimal number of chromatin folding motifs (CFMs) was determined using topic coherence, which measures how consistently contacts within each motif co-occur across the dataset^48^ (Methods). The coherence score reaches a maximum between N_CFM_∼25-30 (**Fig. 1d**, left panel), indicating that the heterogeneous conformational ensemble can be captured by a limited set of recurrent motifs. In parallel, single-allele reconstruction scores increased monotonically with N_CFMs_ (**Fig. S2a**), indicating that additional motifs progressively improve the representation of chromatin traces. Importantly, motifs remained highly similar when varying the number of CFMs around the coherence maximum (N_CFM_=20-30, **Fig. S2b**), supporting stability of the inferred structural patterns. Finally, we assessed over-parameterization by monitoring the emergence of empty CFMs, which are absent below N_CFMs_ = 40, but increased substantially above this threshold (**Fig. 1d**, right panel). Together, these analyses indicate that N_CFM_=30 provides a robust and parsimonious representation of the conformational ensemble, maximizing motif coherence and reconstruction accuracy while avoiding over-parameterization.

Applying 3DTopic to the *Pdx1* dataset revealed a dictionary of recurrent CFMs that capture the dominant patterns of single-allele chromatin organization (**Figs. 1e, S2c–d**). Many motifs correspond to physically interpretable interaction patterns, including contacts between neighboring genomic regions (domain-like motifs, 33%) and contacts between genomically distant loci (loop-like motifs, 57%). A small subset of motifs (10%) capture technical-related patterns and were therefore classified as N/A (**Fig. S2e**). As expected, the structural features emphasized by the decomposition depend on the physical interaction scale defined by the distance cutoff. Lower thresholds (50 nm) produced very sparse contact maps that reduced co-occurrence statistics and led to noisier motifs, whereas higher thresholds (150–200 nm) increased contact density and progressively merged distinct CFMs (**Figs. S3a–c**). The 100-nm threshold therefore provides a robust interaction scale that resolves reproducible motifs at the TAD level while maintaining sufficient contact statistics for reliable inference. All analyses below use these fixed settings (N_CFM_=30; distance cutoff = 100 nm) unless stated otherwise.

To assess the reproducibility of the inferred motifs, we performed over 200 independent trainings and projected the resulting CFMs into a common embedding space using Uniform Manifold Approximation and Projection (UMAP, see *Methods*). CFMs derived from independent decompositions clustered together, indicating that 3DTopic robustly recovers similar conformational motifs across independent trainings (**Fig. 1f**, Methods). Each reference CFM corresponded to a distinct cluster (**Figs. 1e, S2d**) and closely resembled the average motif of the associated cluster (**Fig. 1f**, stars and insets). Direct comparison of motifs obtained from independent decompositions further confirmed the reproducibility of the inferred structural patterns (**Fig. S4a**).

To validate the ability of 3DTopic to reconstruct experimental datasets, we tested how models trained on experimental 3D structures compared to models trained on Gaussian chain polymers, or on randomized contact maps (**Fig. S4b**). As expected, models derived from experimental datasets substantially outperformed these controls and successfully reconstructed the experimental ensemble proximity frequency map (**Fig. S4c**), indicating that the inferred CFMs capture genuine structural patterns present in the data. Importantly, while 3DTopic is primarily designed to identify common patterns in 3D chromatin organization, it also provides a compact and faithful approximation of individual chromatin traces by representing each single-allele contact map as a weighted combination of learned CFMs (*Methods*). We leveraged this property of 3DTopic to show that our reference decomposition faithfully reconstructs single-allele contact maps (**Figs. S4d-e**), with the dominant contacts in each trace largely explained by the most strongly weighted CFMs (**Fig. S4f**). These results indicate that a substantial fraction of single-allele chromatin variability can be captured by combinations of recurrent folding motifs.

### Motif-based organization generalizes across loci and species

To generalize these results, we imaged two additional loci containing disease-relevant genes: *Isl1* and *Ins2*. *Isl1* maintains the beta-cell gene programme via a broad cluster of enhancers^42^ and gene deletion disrupts islet architecture and glucose tolerance^49^, while *Ins2* (and the single human INS orthologue) encodes insulin itself and is driven by promoter/enhancer contacts bound by PDX1 and other β-cell factors, with coding or regulatory mutations leading to hyperglycemia and neonatal diabetes^36,50–52^. As with the *Pdx1* locus, single 3D chromatin traces from the *Isl1* and the *Ins2* loci could be effectively decomposed into loop and domain CFMs (**Figs. 2a-b** and **S5-S6**). Thus, individual 3D chromatin structures of other mouse loci can also be decomposed into sets of motifs representing mainly short-range interaction domains and distal loops.

**Figure 2.**
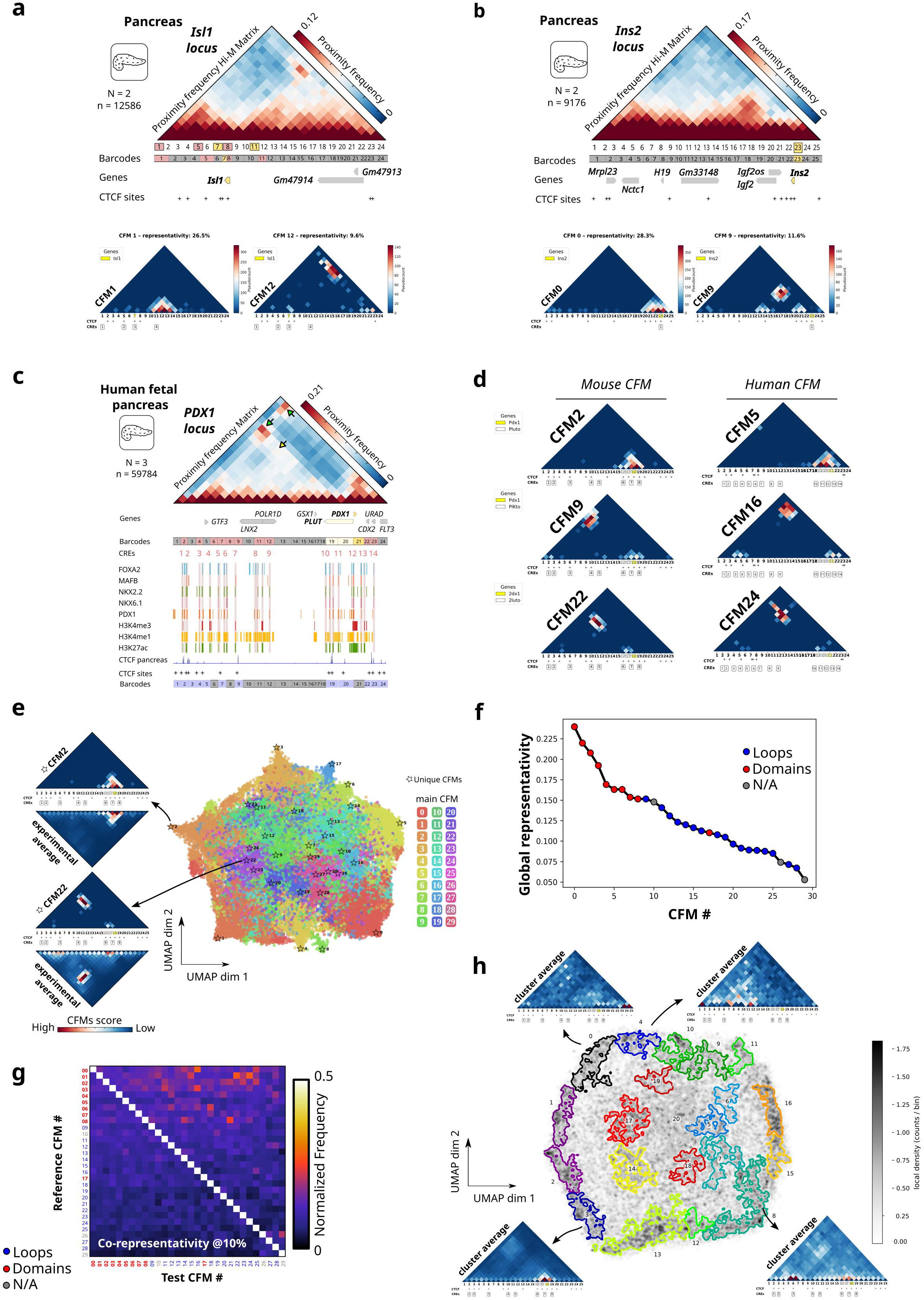
CFMs generalize across diverse mouse genomic loci, show conservation across species, and quantify single-allele co-occurrence patterns. **a-b** Top: Hi-M proximity frequency matrices for the *Isl1* (chr13:116102031-116759941, mm10) (**a**) and the *Ins2* locus (chr7:142520024-142699856, mm10) (**b**) in mouse pancreas. Middle: Hi-M barcodes, genes, and CTCF binding sites. Bottom: subsets of domain and loop CFMs, selected from the complete gallery of 30 CFMs from a single decomposition (see Figs. S5c, S6c, for complete list). **c** Hi-M proximity frequency matrix of the *PDX1* locus (chr13:27902436–28572561, hg19) in human fetal pancreas, shown together with identified CREs, pancreatic TF and CTCF binding sites. **d** Examples of Pdx1-associated CFMs from mice (left) that are also present in the dictionary of human CFMs (right). **e** UMAP of chromatin traces decomposed using the set of CFMs displayed in **Fig. S2d**. Single traces are color-coded by the CFM with the highest weight in the decomposition. Stars indicate the UMAP localization associated with decompositions containing a unique CFM component from the reference decomposition. On the left, two examples showing a reference CFM (top) together with the average contact matrix of all chromatin traces for which this CFM has the highest weight in the decomposition (bottom) (orange dots for CFM2, and violet dots for CFM22). **f** Graph of CFMs representativity for pancreatic cells. Representativity is defined as the frequency at which loop and domain CFMs appear with at least 10% weight in pancreatic cells. Red circles represent domain CFMs, blue circles represent loop CFMs, and gray circles represent CFMs that are not attributed to either loops or domains (N/A). Other representativity thresholds are shown in **Fig. S8a**. **g** Matrix representing the co-representativity (conditional probability) of CFMs occurring in more than 10% in pancreatic cells. The matrix is asymmetric because of the higher frequency of domain over loop CFMs impacting the conditional probabilities. (See *Methods*). **h** UMAP projection of raw binarized contact traces, clustered using HDBSCAN based on local density, with darker regions representing higher-density areas. High-density clusters are primarily observed at the periphery of the embedding. Around the embedding, examples of mean contact maps computed from traces belonging to some cluster are shown.

To assess whether similar folding motifs are identified in other species, we acquired Hi-M datasets at the *PDX1* locus in developing human fetal pancreas (**Figs. 2c, S7a-b**). Applying 3DTopic decomposition to the human dataset identified both domain and loop CFMs (**Figs. S7c-d**). Notably, motifs linking *Pdx1-Plut* to its proximal and distal enhancers in mice (CFM2, 9, 22) were also found in humans (CFM5, 16, 24) (**Fig. 2d**). Consistently, a more thorough comparison of mouse and human CFMs at the *Pdx1/PDX1* locus revealed that over 80 % of mouse CFMs have human counterparts (**Fig. S7e**, and *Cross-species comparison of human and mouse CFMs* in Methods). This motif conservation may originate from the overall similar arrangement of *cis*-regulatory regions and CTCF sites at the *Pdx1/PDX1* regions (**Fig. 2c**), suggesting that these genetic elements underlie the formation of CFMs (see next section). Thus, CMFs underlie chromatin organization at the PDX1 human locus, and similar patterns of 3D organization can be used to describe the conformational space explored by chromatin at this locus in both mouse and human. Together, these results show that motif-based organization provides a general description of single-allele chromatin folding across loci and species, indicating that diverse regulatory regions can be described by combinations of recurrent folding motifs.

### Single-allele chromatin states arise from combinations of motifs with distinct occupancies and co-occurrence

Next, we asked whether decomposition of single traces requires multiple motifs, which motifs are more common, and whether motifs were mutually exclusive or instead occurred concurrently. For this, we mapped the CFM decompositions of individual traces using a UMAP embedding, and color-coded each chromatin trace by the CFM with the highest component (**Fig. 2e**). As expected, chromatin traces dominated by the same reference CFM clustered together and shared its characteristic structural features (**Fig. 2e**). However, we note that most traces in the dataset were decomposed into multiple CFM components (**Fig. S2c**), and were most often positioned in the UMAP between reference CFMs (**Fig. 2e**, stars). Therefore, while many traces display chromatin conformations similar to that of single reference CFMs, their structural reconstruction most often requires the use of multiple motifs.

Next, we quantified motif occupancy in single traces by measuring the fraction of alleles in which each domain or loop CFM contributes at least 10% of the decomposition (**Figs. 2f and S8a-d**). This single-allele frequency of discrete 3D conformations cannot be read directly from ensemble contact maps, which mix heterogeneous conformations into averaged probabilities. Domain CFMs consistently showed the highest representativity (10–25%) across the *Pdx1*, *Isl1* and *Ins2* mouse loci as well as for the human *PDX1* locus (**Figs. 2f, S8a-d**, black lines). In contrast, loop CFMs consistently exhibited lower representativity (5–15%), comparable to looping frequencies between distant CTCF sites measured in live cultured cells (3–30%)^24,25^. As expected, estimated representativity varies with the minimum motif weight threshold (**Figs. S8a-d**, blue, green and yellow lines). Thus, thanks to the ability of 3DTopic to quantify motif usage across single-alleles, we show that short-range domain-like conformations are consistently adopted more frequently than long-range looping states within TADs.

Ensemble methods often display multiple overlapping patterns of interactions within TADs, yet it is unclear whether these features reflect independent conformations in different cells, or instead co-occur within the same single alleles^53,54^. We leveraged the ability of 3DTopic to quantify motif weights in single alleles to discern between these alternative models. Most single traces are composed of mixtures of CFMs rather than a single dominant motif (**Figs. S2c, S5b, S6b, S7c**), indicating that single traces typically contain multiple overlapping structural patterns. To test whether specific motifs preferentially appear together, we computed a co-representativity measure: for each reference CFM, we calculate the conditional probability that a second CFM is also present above a fixed weight threshold in the same allele (10%, Fig. 2g; Methods). We note that the most dominant co-occurance scores correspond to domain-loop CFMs associations (**Fig. 2g**, red bins), consistent with single-allele chromatin structures being commonly composed of both domain-like and loop-like interactions, rather than representing purely domain-only or loop-only states.

To assess whether direct embedding of raw contact maps could recover recurrent structural patterns without prior decomposition, we applied UMAP to binary single-allele contact maps from the *Pdx1* locus (**Fig. 2h**). The resulting embedding organizes traces according to their overall similarity, yielding a continuous representation of the conformational ensemble. To determine whether this representation resolved structural patterns comparable to those identified by 3DTopic, we performed density-based clustering in the UMAP space and averaged raw contact maps within each cluster. While some clusters resembled domain CFMs (e.g. cluster 3), most clusters merged multiple interaction patterns together (e.g. clusters 0, 4, 8) (**Fig. 2h**), preventing the separation of the underlying structural motifs recovered by 3DTopic. In addition, this direct embedding approach does not provide an explicit basis representation of chromatin traces and therefore does not allow assignment of per-trace contributions of recurrent motifs. Together, these results show that single-allele chromatin conformations are structured as combinations of recurrent motifs with distinct occupancies and non-exclusive co-occurrence, a level of organization that is not directly resolved in raw representations of chromatin contacts.

### Simulations and cohesin perturbations provide mechanistic interpretation of CFMs

To shed light into the mechanistic interpretability of CFMs, we first performed molecular dynamics simulations^55^. We generated polymer ensembles under two simplified mechanisms and asked whether 3DTopic recovers domain– and loop-like motifs from the resulting single-polymer conformations. We simulated local interaction–driven polymer conformations using a block copolymer model where a subset of neighboring monomers was allowed to interact when found at close spatial proximity (**Figs. S9a-b**) (see *Methods*)^56^. Using this model, we generated 7500 independent single polymer structures and decomposed them using 3DTopic. Polymer structures could be decomposed by domain-like CFMs encompassing the interacting monomers (**Fig. S9a**). Second, we performed molecular dynamics simulations of loop extrusion with two convergent CTCF sites (**Figs. S9c-d**) (see *Methods*). The main CFM in the decomposition of 7500 single polymer conformations was a loop between CTCF sites (**Fig. S9c**). This motif corresponds to conformations where loop extruding factors (LEFs) stall at convergent CTCF sites, while the remaining motifs represent incompletely extruded polymers. Thus, 3DTopic recovers domain-like and loop-like CFMs from block copolymer and loop extrusion simulations.

Next, we wondered if this simple *in silico* approach would be able to reconstruct the experimentally observed CFMs. For this, we first performed block copolymer simulations where mouse pancreas accessible regions along the *Pdx1* locus were used as proxies for *cis*-regulatory elements, and allowed to interact with each other when in close spatial proximity (**Fig. 3a**). As expected, 3DTopic decompositions produced only domain CFMs (**Figs. 3a, S9e**). Increasing the interaction strength between monomers led to complex CFMs with multiple short– and long-range interactions, and no discrete loop-like CFM was observed (data not shown). Thus, block co-polymer simulations based on interactions between accessible chromatin at the *Pdx1* locus recapitulate predominantly short-range interaction patterns described by domain CFMs.

**Figure 3.**
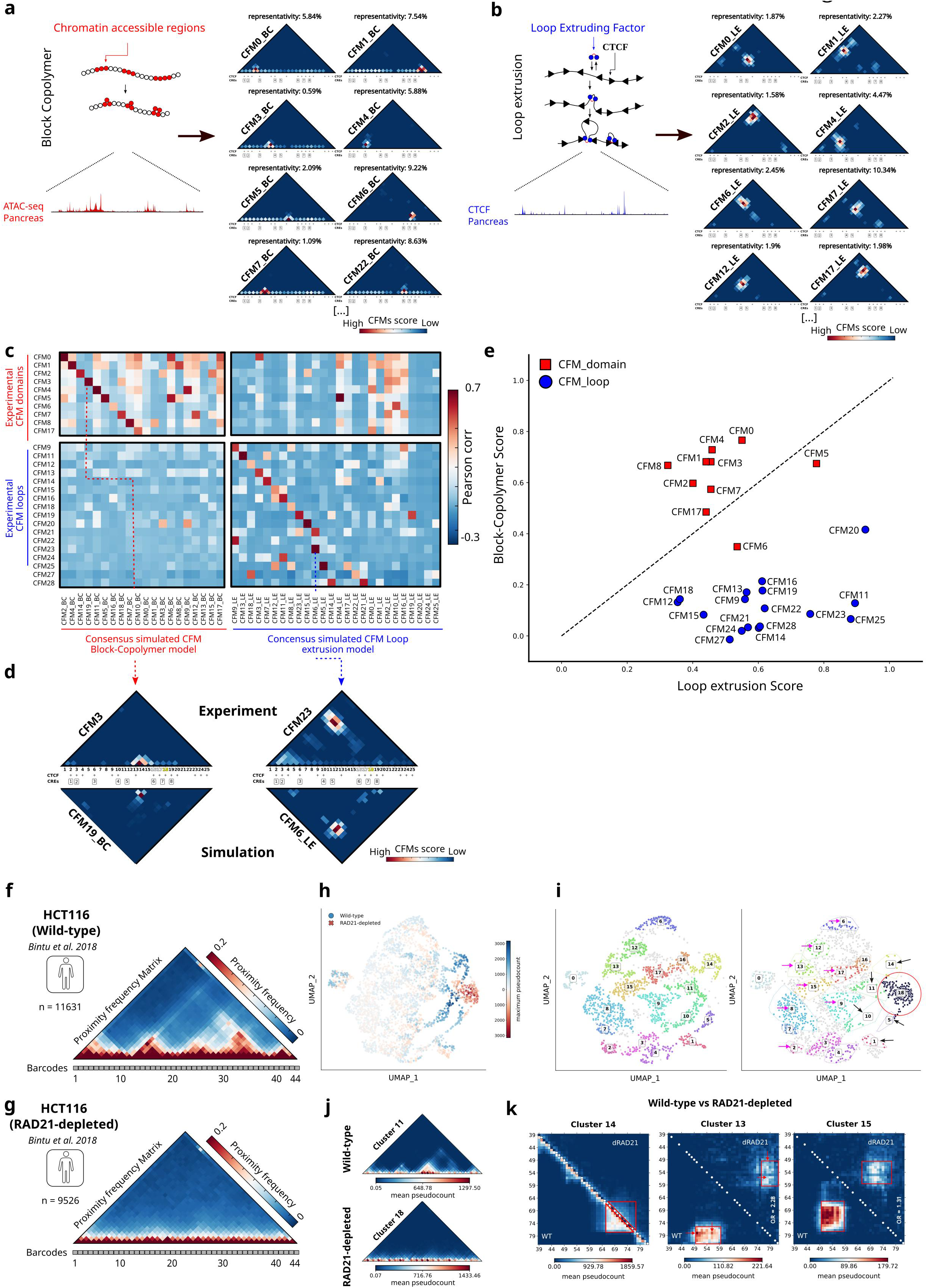
Polymer simulations and acute RAD21 depletion reveal major role of loop extrusion in CFM formation. **a** Subsets of CFMs obtained from a block copolymer simulation of the *Pdx1* locus using chromatin accessible regions from ATAC-seq data (C. Liu et al. 2019) (see *Polymer simulations* in *Methods*). Complete gallery of CFMs are provided in **Fig. S9e**. **b** Subsets of CFMs from a loop extrusion model of the *Pdx1* locus using CTCF sites from CUT&Tag data (R. R. Wang et al. 2022) (see *Polymer simulations* in *Methods)*. A complete gallery of CFMs is provided in **Fig. S9f**. **c** Cross-correlation matrix comparing the experimental CFMs of the reference decomposition (**Fig. S2d**) with consensus CFMs from either a block copolymer model or a loop extrusion model (**Figs. S9e-f**, and *Comparison of experimental and simulated CFMs* in *Methods*). **d** Examples of experimental (top) and simulated CFMs (bottom). **e** Graph representing the block-copolymer score (y-axis) versus the loop extrusion score (x-axis) for each individual experimental CFM from the *Pdx1* locus. Domain CFMs are represented by red squares, while loop CFMs are depicted by blue circles. **f-g** Proximity frequency matrices for the locus traced in Bintu *et al.*^18^ using HCT116 cells for either (**f**) wild-type or (**g**) RAD21-depleted conditions. Barcodes are displayed as grey boxes below the maps. **h** Embedding of CFMs pooled from 100 runs of 3DTopic decomposition from wild-type (total 2000 components) and RAD21-depleted (total 1500 components). Each CFM is colored by its maximum pseudocount, which reflects the prevalence of the corresponding conformational motif within the training dataset (blue: enriched in wild-type; red: enriched in RAD21-depleted). **i** Clustering of CFMs in the embedded space. Clusters identified in wild-type cells (left) are overlaid as hulls on the CFM embedding derived from RAD21-depleted cells (right). The red circle highlights the main cluster in RAD21-depleted cells, black arrows point to clusters detected in the WT conditions and absent in RAD21-depleted cells (comprising domains), and pink arrows indicate hulls enclosing CFMs in RAD21-depleted cells that form clusters in WT cells, i.e., sets of CFMs common to both conditions (comprising mostly loop– and stripe-like structures). **j** Average representations of cluster 11 (wild-type) and cluster 18 (RAD21-depleted). **k** Side-by-side comparison of average cluster CFMs for wild-type (bottom left) and RAD21-depleted (top right) data in three different clusters (clusters 14, 13, and 15). Red boxes and arrows represent the main pattern of interaction represented by a CFM, while the white circles represent CTCF binding sites. The odds ratio of the long-range pattern between two CTCF binding sites (see Methods) is indicated on the right edge of relevant maps.

To assess the contribution of loop extrusion to CFM formation, we performed loop extrusion simulations where CTCF sites were positioned using the CTCF binding sites from mouse pancreas at the *Pdx1* locus (**Fig. 1b**). In this case, most reconstructed motifs represented loops (**Figs. 3b, S9f**). In addition, domain CFMs were also observed when CTCF sites were located in close genomic proximity (**Figs. S9f-g**). Therefore, loop extrusion simulations of the *Pdx1* locus can reproduce loop CFMs and a subset of domain CFMs.

To determine whether these mechanisms can reproduce the experimentally observed CFMs, we compared simulated and experimental motifs using Pearson correlation analysis. Notably, most of the experimentally obtained domain CFMs can be matched by at least one CFM simulated using a block copolymer model (**Fig. 3c**, top-left quadrant, and **Fig. 3d**). We remark that block copolymer simulations are unable to recapitulate experimental loop CFMs under the tested parameter regime with weak monomer interactions (**Fig. 3c**, bottom-left quadrant). Instead, these CFMs can be properly restored by loop extrusion simulations (**Fig. 3c**, bottom-right quadrant, and **Fig. 3d**). In addition, we note that loop extrusion can also reproduce a subset of the experimental domain CFMs that arise from loops between genomically-close CTCF sites (**Fig. 3c**, top-right quadrant). In agreement with these analyses, most loop CFMs exhibited low block copolymer scores (<0.2) and high loop extrusion scores (>0.5) (**Fig. 3e**, see *Methods*). In contrast, domain CFMs display in general a better agreement with block copolymer CFMs (e.g. CFM0-4, CFM7-8), yet several domain CFMs can be equally or better matched to loop extrusion CFMs (e.g. CFM5-6), consistent with the idea that domains can also be formed by proximal CTCF sites (**Fig. S9g**). Thus, most experimental CFMs can be reproduced by loop extrusion simulations with block copolymer models being required to recover a subset of domain CFMs.

To test whether loop extrusion contributes to the formation of chromatin folding motifs *in vivo*, we reanalyzed published chromatin tracing data from a mammalian system in which RAD21, an essential cohesin subunit, was degraded^18^. As expected, proximity frequency maps from wild-type and RAD21-depleted cells revealed a marked reduction in both short– and long-range contacts following cohesin loss (**Figs. 3f–g, S10a-b**). To quantify these effects on chromatin folding motifs, we first binarized the pairwise distance matrices into contact matrices using the optimal distance cutoff of 150 nm (selected as described previously, **Figs. S10c-d**), and then selected the optimal number of CFMs used for 3DTopic decompositions (N_CFMs_=20, **Fig. S10e**). Next, we performed 100 independent 3DTopic decompositions for both conditions and embedded the resulting CFMs into a common UMAP (**Fig. 3h**). CFMs from wild-type and RAD21-depleted cells occupied distinct regions of the embedding, indicating that loop extrusion contributes to defining different conformational ensembles.

Clustering analysis of wild-type CFMs revealed that several of the most populated clusters (1, 5, 10, 11, 14) were strongly underpopulated in RAD21-depleted samples (**Fig. 3i**, black arrows). These major wild-type CFMs correspond to domain-like CFMs and show a pronounced reduction after RAD21 degradation (**Figs. 3j** top panel, and **S10f**). Conversely, the cluster most enriched in RAD21-depleted alleles (cluster 18, **Figs. 3h, j**), not detected in the wild-type condition, represents polymer-like configurations (**Fig. 3j**, bottom panel). Thus, RAD21 degradation leads to the loss of domain-like CFMs and the appearance of polymer-like motifs.

To determine how loop-like CFMs were affected by RAD21 depletion, we examined wild-type CFM clusters displaying long-range interaction patterns. Most of these clusters were significantly underpopulated in RAD21-depleted cells (**Fig. 3i**, pink arrows) where they displayed a pronounced reduction in pseudocount / mean CFM score (**Figs. 3k** and **S10f**), indicating that RAD21 degradation leads also to the loss of the long-range interaction patterns represented by loop CFMs. Together, these analyses provide a mechanistic interpretation of CFMs, with cohesin-mediated loop extrusion playing a major role in generating loop– and domain-like motifs, and local *cis*-regulatory interactions instead contributing to the formation of a subset of domain-like motifs.

### Tissue specific folding arises from differential usage of a common motif repertoire

Cell-type-specific transcription is regulated in large part by *cis*-regulatory elements that can drive changes in 3D chromatin organization between tissues^14^. However, ensemble methods are unable to discern whether *cis*-regulatory rewiring between tissues modifies the repertoire of 3D motifs, or instead modulates the usage of a single set of common motifs. We leveraged the ability of 3DTopic to decompose single-allele conformations into a set of interpretable motifs to test between these alternative models. The *Pdx1* locus provides a useful paradigm, as it harbors multiple genes with distinct expression profiles: for instance, *Pdx1* is expressed primarily in the pancreas, *Gsx1* in the brain, and *Urad* in the liver, while other genes such as *Polr1d*, *Pan3*, *Gtf3*, or *Mtif3* are expressed more broadly (**Figs. 4a, S11a**). Most CTCF binding sites in the *Pdx1* locus are shared across tissues, with only a single site absent from pancreas at the resolution of our library (**Fig. 4a**, barcode 22). In contrast, ATAC-seq peaks vary more extensively, with 17 out of 38 peaks absent in the pancreas but present in at least one other tissue. Yet, at the resolution of our library, only 5 out of 25 barcodes (11, 14, 21, 22, 23) harbor such tissue-specific accessibility sites (**Fig. 4a**). We therefore asked whether tissue-specific accessibility differences underpin changes in the motifs themselves or instead in motif usage. For this, we first generated Hi-M data from six additional tissues: brain, kidney, liver, lung, lymph node and thymus. The resulting proximity and PWD distance Hi-M maps revealed noticeable structural variation within the *Pdx1* locus between tissues at the ensemble level (**Figs. 4b, S12a**). Consistently, other loci (e.g. *Ins2*) also displayed 3D structural differences across tissues (**Figs. S12b-c**).

**Figure 4.**
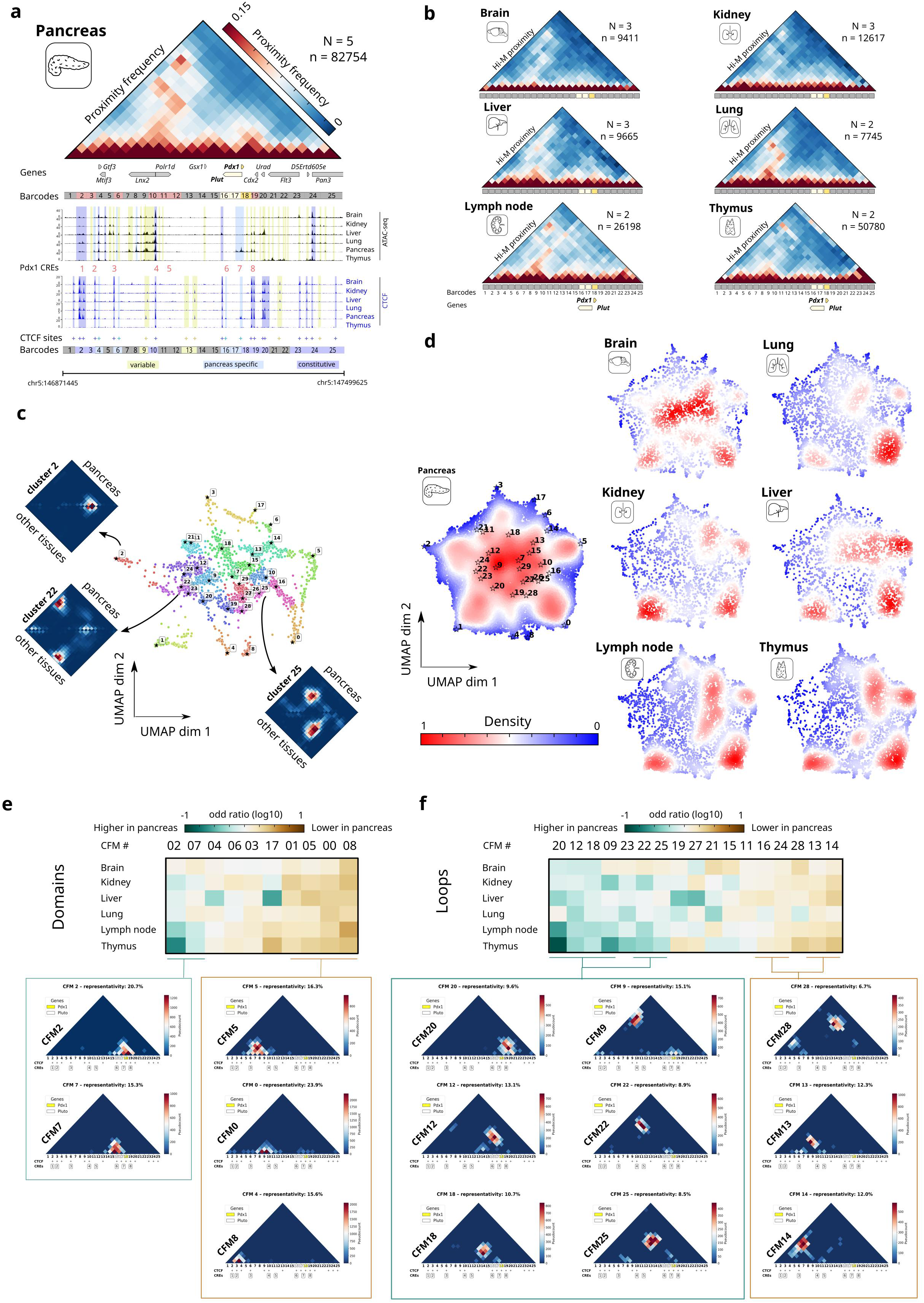
Single cell structures from seven mouse tissues can be decomposed with the same set of CFMs. **a** Top: Hi-M proximity matrices for the *Pdx1* locus in the pancreas. *N* represents the number of biological replicates, and *n* the number of traces. Bottom: ATAC-seq (black) and CTCF ChIP-seq (Brain, Kidney,Liver,Lung,Thymus) or CUT&Tag (Pancreas) (blue) profiles (see *Methods*) along the Pdx1 locus in six different tissues, with gene locations and CTCF binding sites. CTCF-binding sites, indicated by “+” signs and Hi-M barcodes, are color-coded as variable (yellow), pancreas-specific (blue), or constitutive (violet) across all examined tissues. **b** Hi-M proximity matrices of the *Pdx1* locus in six additional mouse tissues: Brain, Kidney, Liver, Lung, Lymph node and Thymus. **c** UMAP embedding of 200 independent trainings using traces from the *Pdx1* locus from Brain, Kidney, Liver, Lung, Lymph node and Thymus. Colors represent different clusters from HDBscan. Stars shows the CFMs originating from the reference training used for the *Pdx1* LR library (shown in **Fig. S2d**). Surrounding the embedding are the cluster-average CFM matrices (computed from all CFMs within each cluster), based on decompositions performed either using pancreas cells only (top; pancreas) or using all non-pancreatic cells (bottom; other tissues). **d** Density distribution of pancreatic cells (left) and six other mouse tissues (right) projected onto the UMAP landscape. Red and blue indicate regions of high and low cell density, respectively. **e-f** OR matrix for domain (**e**) and loop (**f**) CFMs taking the pancreas as reference tissue. Green indicates OR < 1 (CFM enriched in pancreas), and orange indicates OR > 1 (CFM enriched in other tissues). Examples of CFMs enriched in pancreas (left) or in other tissues (right) are displayed below the OR matrices.

Next, we asked whether the structural patterns observed in different tissues were distinct from, or instead shared with, those mapped in pancreas. To address this, we trained CFMs exclusively on data from non-pancreatic tissues and jointly embedded these with pancreas-derived CFMs into a common low-dimensional representation of the motif space (**Fig. 4c**). We then identified recurrent conformations using HDBSCAN clustering (see *Methods*) to group similar CFMs across independent decompositions. Notably, reference pancreas CFMs localized to clusters defined by CFMs from other tissues, and direct one-to-one comparisons revealed that pancreas CFMs and the average CFMs of corresponding clusters were nearly identical (**Figs. 4c, S13a**). Thus, despite differences in tissue-specific accessibility patterns, the conformational motifs identified in pancreas are the same as those obtained when combining chromatin traces from multiple other tissues and are sufficient to reconstitute the 3D structure of the *Pdx1* locus with high accuracy (**Figs. S13b-c**). Together, these results support a model in which *cis*-regulatory interactions act by changing the usage of motifs between tissues rather than by creating tissue-specific *de novo* motifs.

To further explore this idea, we decomposed chromatin traces from other tissues using the reference CFM dictionary derived from pancreas and visualized their distributions in UMAP space. Projection of all tissues into the same reference embedding revealed extensive tissue-dependent shifts in density for both the *Pdx1* locus (**Fig. 4d**) and the *Ins2* locus (**Fig. S13d**). Thus, while different tissues display distinct frequencies of CFM usage, they do not generate novel architectural motifs at the loci examined. Instead, tissue-specific *cis*-regulatory logic acts by modulating the relative prevalence of loop and domain CFMs within a shared motif repertoire.

*Pdx1* is specifically expressed in the pancreas (**Fig. S11a**), therefore we expect CFMs linking *Pdx1* to its proximal and distant enhancers (CFMs 2,7 and CFMs 9, 18, 20, 22, 25) to shift their usage between pancreas and other tissues. To test this hypothesis, we performed a statistical analysis using the Odds Ratio (OR, *Methods*). An OR > 1 indicates higher usage in the test tissue (e.g. brain, kidney, etc), whereas an OR < 1 indicates higher usage in pancreas; we considered |log10(OR)| > 0.1 as significant (**Fig. S13e**). Notably, domain CFMs linking *Pdx1* to its proximal enhancers were significantly overrepresented in pancreas (**Fig. 4e**). Similarly, loop CFMs involving *Pdx1* and its distant enhancers were consistently more predominant in pancreas than in other tissues (**Fig. 4f**). We note that most *Pdx1*-associated loop CFMs also contained CTCF anchors specifically accessible in pancreas (barcodes 4, 6, 16, 17), therefore their higher frequencies in pancreas are consistent with contributions from pancreas-specific CRE interactions and/or pancreas-specific CTCF loops. Overall, these results show that despite large differences in transcription and *cis*-regulatory activity, chromatin traces from different tissues can be described using the same motif repertoire, with tissue-specific regulation reflected in changes in the frequency of motif usage.

### Transcriptional activation reshapes chromatin folding through motif-specific changes in single-allele subpopulations

To reveal how transcriptional activation reshapes single-allele chromatin conformations, we quantified changes in CFM usage between closely related pancreatic cell types where *Pdx1* is differentially transcribed. For this, we performed sequential RNA-FISH imaging in the mouse pancreas followed by Hi-M at the *Pdx1* locus (**Figs. 5a-b**). We classified pancreatic cells into beta cells, which express insulin (Ins1/Ins2), or exocrine cells which do not express any of the major endocrine cell markers, i.e. Ins1/Ins2, glucagon (*Gcg*), and somatostatin (*Sst*) (**Fig. 5b**, top). Next, we reconstructed ensemble Hi-M maps for beta and exocrine cells (**Fig. 5b**, bottom), and used them to compute a differential pairwise distance map displaying regions that are, on average, closer in beta or in exocrine cells (**Fig. 5c**).

**Figure 5.**
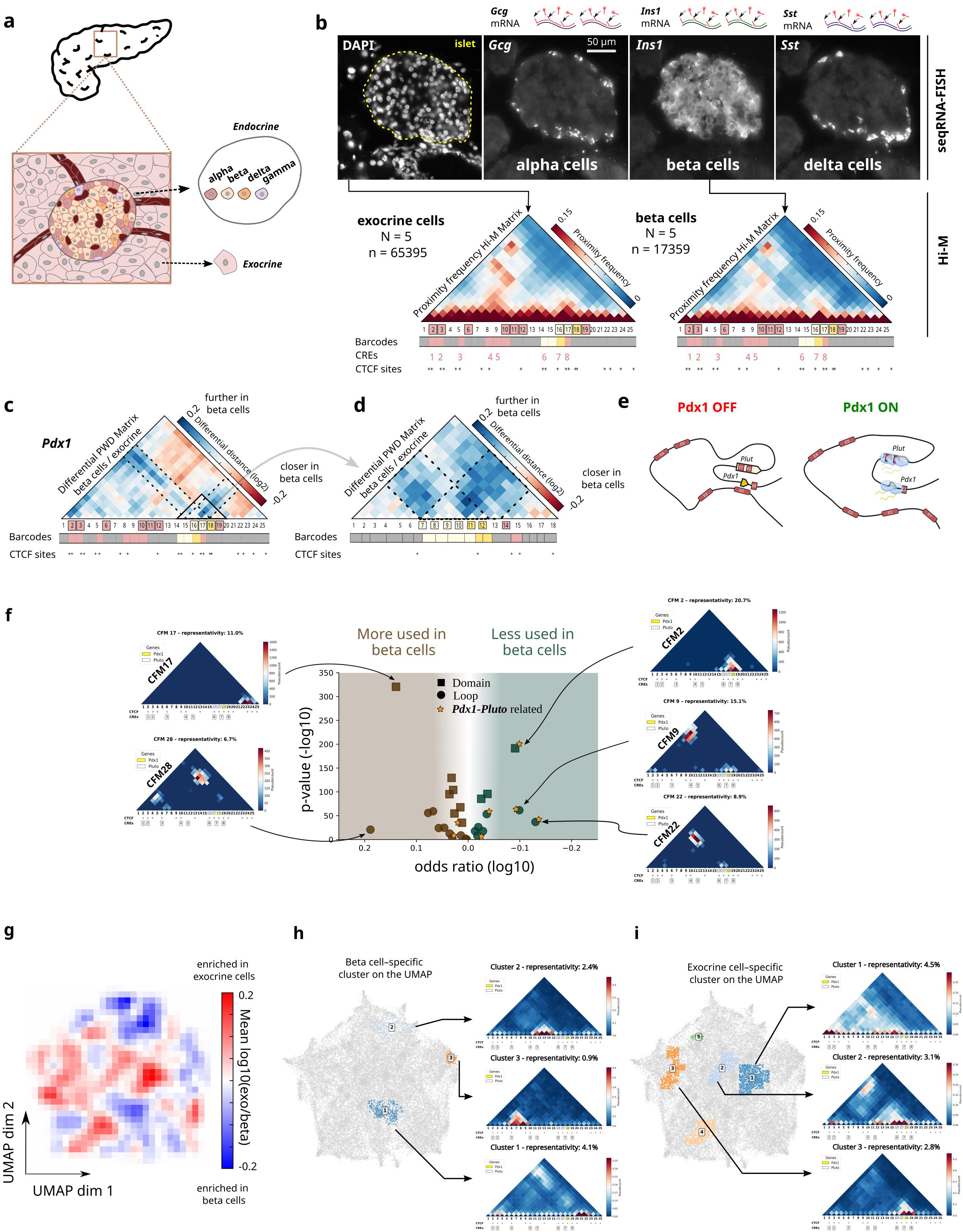
Cell type specific 3D chromatin organization can be described by differential usage of CFMs. **a** Schematic illustrating the different cell types of the pancreas. The pancreas is divided into exocrine and endocrine tissues, with the islet representing the basic unit of the endocrine tissue. Islets are constituted of various cell types, including alpha, beta, delta, and gamma cells. **b** Top: Grayscale maximum intensity projection of a DAPI-stained nuclei image, centered on a pancreatic islet. Exocrine cells (outside the contour in dashed lines) are defined as cells not displaying any of the endocrine markers (*Ins1/Ins2, Gcg, Sst*). Grayscale maximum intensity projection of *Ins1*, *Gcg* and *Sst* RNA-FISH markers, corresponding to alpha, beta, and delta-cells, respectively. Beta cells are defined as *Ins1+* cells. Bottom: Corresponding proximity frequency matrices for exocrine (left) and beta (right) cells along the *Pdx1* locus. *N* represents the number of replicates, *n* represents the number of traces. CTCF sites are shown as “+” signs, CREs are shown in pink and *Plut* and *Pdx1* are represented by white and yellow rectangles, respectively. **c-d** Differential PWD Hi-M matrices between beta cells and exocrine cells. Blue and red represent higher and lower distances in beta cells compared to exocrine cells, respectively. Solid triangle represents the region visualized in panel (**d**) at higher resolution (see Pdx1 HR in *Methods*). Dashed lines represent the bins occupied by *Pdx1* and *Plut*. **e** Schematic representing the main results from analysis of the ensemble differential distance maps, where *Pdx1-Plut* becomes more distant to all other barcodes in the locus in beta cells where it is expressed. **f** Volcano plot showing the log10 odd ratio (x-axis) between exocrine cells and beta cells, and –log10 (P-value) for each individual CFMs from 500 different decompositions. Domain CFMs are represented by squares, while loop CFMs are depicted by circles. Stars indicate CFMs related to the *Pdx1* gene. **g** Differential UMAP density plot showing the relative density of exocrine and beta cells in the UMAP space. Blue indicates higher density, and red indicates lower density in beta cells compared to exocrine tissue. **h** UMAP highlighting subpopulations of traces overrepresented in beta cells. Examples of averaged proximity maps for each subpopulation are shown. **i** Same as panel (**h**), but showing subpopulations of traces overrepresented in exocrine cells.

Surprisingly, this differential map shows that *Pdx1-Plut* are on average farther away from their proximal and distal enhancers and from all other loci in their TAD (**Fig. 5c**), specifically in beta cells where they are highly differentially expressed (**Fig. S14a**). To further validate this result, we performed Hi-M using a high-resolution oligopaint library that zooms-in on the *Pdx1-Plut* region, which further revealed a local decompaction of the *Pdx1-Plut* gene unit in beta cells (**Figs. 5d, S14b-c**). Finally, we also observed local gene decompaction for *Isl1*, an endocrine cell-specific TF expressed specifically in beta and not in exocrine cells (**Fig. S14d-f**)^49,57^. In brief, these results indicate that higher transcriptional activity is accompanied by an increase in average distance between the promoter and other regions within its TAD (**Fig. 5e**). However, ensemble differential maps do not reveal whether this arises from loss of short– or long-range loops, or both, nor whether these changes reflect a coordinated rewiring across all alleles or from specific subpopulations of single-allele conformations.

To tackle these questions, we first analyzed the relative changes and variability in CFM utilization between chromatin traces from beta and exocrine cells. For this, we plotted the OR versus the p-value for each CFM (**Fig. 5f**, *Methods*). Remarkably, CFMs encoding interactions involving *Pdx1-Plut* with proximal and distal regions (CFM2, CFM9, CFM22, **Fig. 5f**) were significantly depleted in beta cells, where *Pdx1* is more highly expressed. CFM2 represents contacts between *Pdx1-Plut* and their proximal CREs, while CFM9 and CFM22 anchor *Pdx1-Plut* to a cluster of distal CREs (CRE1 and CRE2). Reconstruction of these CFMs using simulations (**Figs. 3c-e**) indicate that they arise from either block-copolymer interactions between neighboring open chromatin regions (CFM2) or from loop extrusion (CFM9, 22) (**Figs. 3c-e**). The depletion of these CFMs is consistent with increased pairwise distances between *Pdx1-Plut* and their proximal and distal enhancers (**Fig. S14g**). Similar conclusions can be reached from high-resolution tracing data of the *Pdx1-Plut* locus and from the *Isl1* locus (**Figs. S14h-k**). Thus, the ensemble-average decompaction of the *Pdx1* locus in beta cells involves depletion of both short– and long-range CFMs linking *Pdx1-Plut* to its enhancers.

These analyses identify which CFMs change between beta and exocrine cells, but do not reveal whether the observed motif shifts reflect diffuse reweighting across most alleles or preferential changes in discrete conformational subpopulations. Because 3DTopic represents each allele as a mixture of CFMs, we can directly discern between these two scenarios. For this, we decomposed single allele conformations from beta and exocrine cells using the reference dictionary of CFMs, embedded these decompositions using UMAPs, and calculated the differential UMAP to estimate subpopulations of traces preferentially over– or under-represented in beta and exocrine cells (**Fig. 5g**). The differences between the UMAPs of these well-defined cell types were considerably smaller than those previously observed between cell types from different tissues (**Figs. 4c-d**), which is consistent with both beta and exocrine cells originating from a common multipotent pancreatic progenitor. Nonetheless, analysis of the differential UMAP reveals clusters specifically enriched in beta (**Fig. 5h**) or in exocrine cells (**Fig. 5i**).

We observe three independent subpopulations of single traces enriched in beta cells (**Fig. 5h**), accounting for the regions of the differential ensemble distance map where distances are smaller in beta cells (**Fig. 5c**). More importantly, five subpopulations are depleted in beta cells and enriched in exocrine cells (**Fig. 5i**). These beta-depleted clusters are dominated by conformations that involve the *Pdx1–Plut* unit itself (cluster 3) and its long-range contacts with CRE3 (cluster 1) or CRE1–2 (cluster 2), and correspond to the enhancer-linking motifs identified above as depleted in beta cells (CFM2, CFM9 and CFM22; **Fig. 5f**). Thus, the ensemble-average loss of enhancer proximity is driven by selective depletion of discrete single-allele conformational states, rather than uniform weakening of all contacts across alleles. Overall, these analyses show that cell-type-specific transcription is accompanied not by a single but by multiple, separable motif-level changes affecting both local and long-range promoter-CRE contacts in distinct subpopulations of cells.

### Obesity/T2D remodels beta-cell chromatin folding in a subset of alleles via motif reweighting

Our previous analyses showed that ensemble differences in chromatin organization of related cell-types can arise from selective reweighting of folding motifs in distinct subpopulations of alleles. Next, we investigated whether similar changes occur during the induction of obesity and T2D, which are characterized by aberrant beta cell transcription of *Pdx1* and other markers. For this, we fed mice with either a normal diet (ND) or a high-fat diet (HFD) for 14 weeks, a well-established preclinical model of obesity and T2D^58^ (**Fig. 6a**). As expected, islet size and number of beta cells per islet increased considerably in HFD-fed with respect to ND-fed mice (**Fig. 6b**)^59^ as well as body weight and blood glucose levels (**Figs. S15a-d**). As previously shown, HFD leads to reduced levels of multiple beta-cell TFs, such as MAFA, PDX1, NKX6.1 and NKX2.2^60–67^ (**Fig. S15e**) which bind to enhancers at the *Pdx1* locus (**Fig. 1b**). In our model, CTCF levels remained unchanged (**Figs. S15f-g**), suggesting that changes in 3D chromatin architecture in beta cells are driven primarily by down-regulation of beta-cell TFs during metabolic stress.

**Figure 6.**
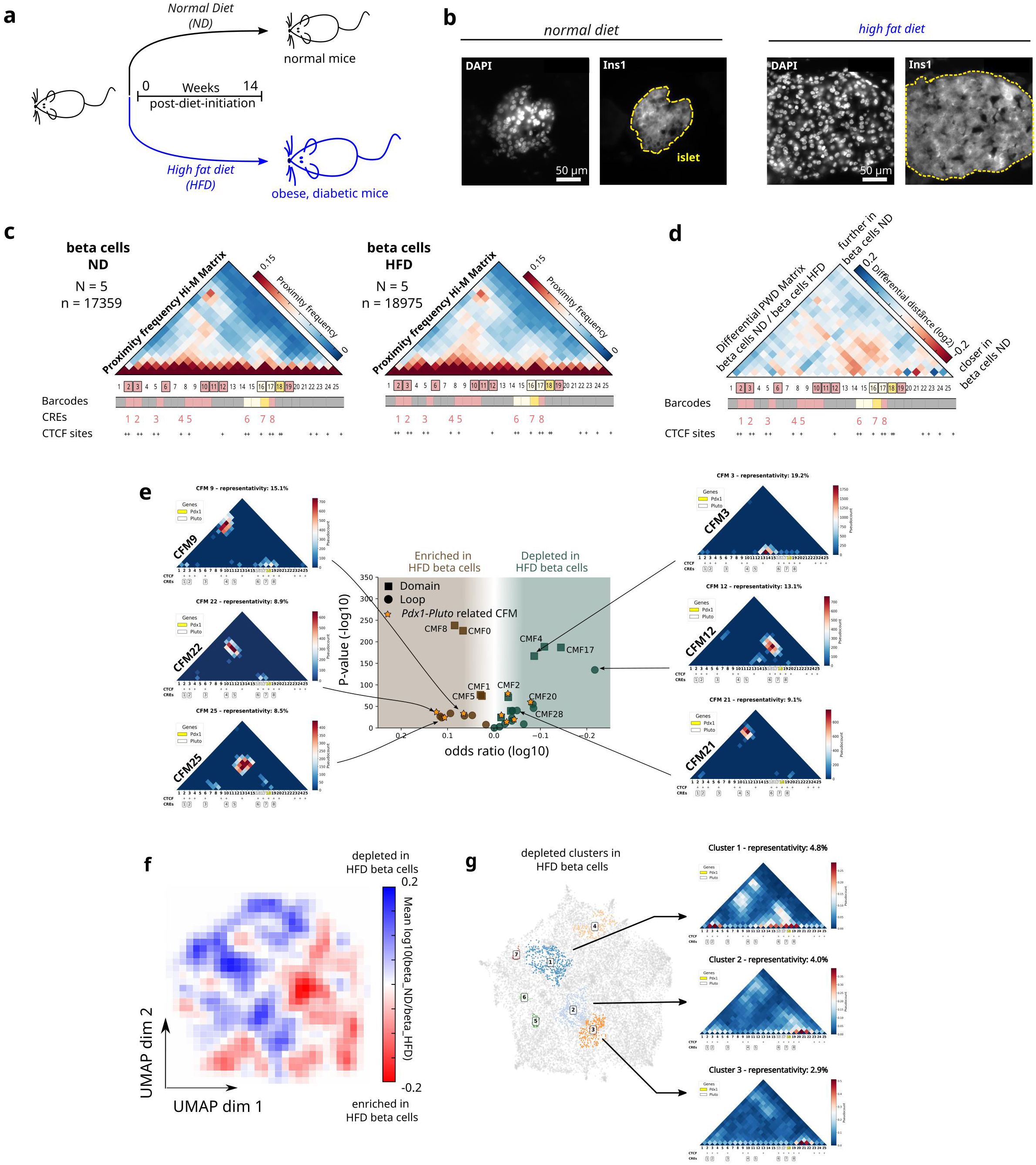
Perturbation of transcription factors and CTCF levels changes CFM utilization. **a** Schematic illustrating the perturbation model to induce type II diabetes (T2D). **b** Left: Grayscale DAPI-stained nuclei image of a mouse pancreas fed with normal diet (ND), centered on a pancreatic islet. Maximum intensity projection of the fluorescence signal from a beta cell marker (Ins1). Right: Microscopy images of mouse pancreas fed with a high fat diet (HFD). In both cases, islets are circled with yellow dashed lines. **c** Proximity frequency matrices along the *Pdx1* locus in beta cells under ND (left) and HFD (right) conditions. **d** Differential PWD Hi-M matrices between beta cells ND and beta cells HFD. Blue and red represent higher and lower distances in beta cells ND compared to beta cells HFD, respectively. **e** Volcano plot showing the log10 odd ratio (x-axis) between beta cells ND and beta cells HFD, and –log10 (p-value) for each individual CFMs across 500 different decompositions. Domain CFMs are represented by squares, while loop CFMs are depicted by circles. Stars indicate CFMs related to the *Pdx1* gene. **f** Differential UMAP density plot showing the relative density of beta cells under ND and HFD conditions. Red indicates enrichment in HFD beta cells (depletion in ND beta cells), while blue indicates depletion in HFD beta cells (enrichment in ND beta cells). **g** UMAP highlighting subpopulations of traces overrepresented in ND beta cells. Examples of averaged proximity maps for each subpopulation are shown. We note that these subpopulations can be decomposed by CFMs enriched in ND beta cells. Cluster 1: CFMs 3, 12 and 21; Cluster 2: CFMs 4 and 20; Cluster 3: CFM4.

In the ensemble, the *Pdx1* locus exhibited broadly similar proximity patterns in beta cells from HFD and ND conditions (**Figs. 6c, S15h**), with only modest shifts apparent in the difference map (**Fig. 6d**). Because *Pdx1* transcription decreases in HFD beta cells as well as in exocrine cells relative to ND beta cells (**Figs. S14a, S15e**), we tested whether metabolic stress induces the same motif-level changes at the *Pdx1* locus as those associated with low *Pdx1* transcription in exocrine cells.

Notably, the ensemble difference map indicates that only a subset of enhancers (CRE1-2) become closer to *Pdx1* in HFD beta cells with the remainder showing only minor shifts (**Figs. 6d, S16a**), in contrast to the uniform reduction in *Pdx1*-enhancer distances observed in exocrine cells (**Figs. 5c, S14g**). Similarly, the pattern of changes in promoter-enhancer distances at the *Isl1* locus differ when comparing ND beta cells to HFD beta cells (**Figs. S16b-d**) or to exocrine cells (**Figs. S14e-f,k**). In short, induction of obesity and T2D leads to beta cell-specific remodeling of enhancer-promoter distances, but these alterations are distinct from those occurring between beta and exocrine cells in ND mice.

To identify which patterns in single-allele 3D chromatin organization are altered in HFD mice, we decomposed beta cell chromatin traces from ND and HFD mice using the reference dictionary of CFMs. We observed significant changes in the usage of both domain and loop CFMs (**Fig. 6e**). Loop CFMs that bring *Pdx1-Plut* spatially close to distal enhancers (CFM9 to CRE1-2, CFM22 to CRE3 and CFM25 to CRE4-5) were enriched in HFD beta cells, mirroring the trend in the beta-to-exocrine comparison. Conversely, CFMs linking *Pdx1* to proximal enhancers (e.g. CFM2) were instead depleted in HFD beta cells. Similarly, CFM shifts between HFD and ND beta cells at the *Isl1* locus (**Fig. S16e)** recapitulated CFM shifts between ND beta and exocrine cells only partially (**Fig. S14j**). Thus, HFD beta cells display CFM shifts that are consistent with lower *Pdx1* transcription, yet distinct from the shift patterns of exocrine cells, suggesting that HFD beta cells do not simply adopt an exocrine-like folding state, even when *Pdx1* transcription is reduced in both contexts.

To test if these motif shifts are confined to defined subpopulations, we embedded ND and HFD beta traces in a common UMAP space and computed a differential density map (**Fig. 6f**). Cluster detection identified only three regions (clusters 1-3) significantly significantly depleted in HFD cells, and with high representativity (**Fig. 6g**). These clusters represented only a small proportion of traces (∼10%), therefore most alleles displayed similar conformations in ND and HFD mice. Notably, clusters 1-3 represent different subpopulations of traces than those depleted in exocrine cells (**Fig. 5h**), further highlighting that HFD beta cells explore a conformational space different from both ND beta and exocrine cells. Notably, the new clusters correspond to subpopulations in which proximal and distal *Pdx1* enhancers (CRE8, CRE1–2, CRE5–6) cluster together within the same chromatin trace while excluding the *Pdx1-Plut* transcriptional unit (**Fig. 6g**). These interaction patterns closely resemble CFM3, CFM12, and CFM21, which are likewise significantly depleted in HFD beta cells (**Fig. 6e**). Together, these results show that the conformational space explored by the *Pdx1* locus is selectively remodelled between HFD and ND beta cells. The most notable conformational changes reflect enrichment in loops between *Pdx1-Plut* and its distant enhancers, and depletion of cell subpopulations where distant enhancers interact with each other and exclude *Pdx1-Plut*. Thus, disease-associated changes in 3D genome organization arise from alterations in motif usage across specific subpopulations of single alleles that remain largely obscured at the level of ensemble maps.

## Discussion

Here, we combine high-resolution multiplexed DNA-FISH imaging with an unsupervised machine-learning framework (3DTopic) to reveal a compact and interpretable representation of chromatin conformational heterogeneity within TADs. Despite the large structural variability observed between single alleles, we show that chromatin conformations can be described as combinations of a limited set of recurrent chromatin folding motifs. This motif-based representation enables direct measurement of motif usage, co-occurrence, and subpopulation structure across tissues, cell types and disease states, providing insight into the following questions:

First, how are TADs organized in single cells? Pioneering ensemble sequencing studies showed that TADs contain multiple overlapping interaction patterns linking CTCF sites, enhancers, and promoters^53,54,68^. Considerable efforts have been devoted to characterizing these substructures^69–72^, yet their organization at the level of individual alleles has remained difficult to resolve. Imaging studies identified higher-order “nano-domains” within TADs^21^, but the lack of genomic specificity limited identification of their molecular identity and underlying mechanisms. Our results show that the heterogeneous conformations explored by single alleles within TADs arise from combinations of a small set of recurrent chromatin folding motifs. These motifs correspond primarily to short-range domain-like interactions and long-range contacts between distal regulatory elements. These findings reconcile ensemble and single-cell views of TAD organization by showing that overlapping interaction patterns arise from combinations of recurrent conformations explored by individual alleles, providing a molecular interpretation for the nano-domain structures observed in imaging studies.

Second, how do loop extrusion and *cis*-regulatory interactions affect 3D chromatin organization within single TADs? Recent ensemble micro-C experiments and theoretical modeling showed that both loop extrusion and transcriptional activity contribute to the regulation of loops within TADs^73,74^. However, the absolute frequencies at which these loops occur in single cells, whether they are mutually exclusive, and how they vary within tissues remain unknown. By quantifying motif frequencies and their co-occurrence at the single-allele level, 3DTopic enables direct comparison between mechanistic models and experimental data. Our analyses reveal that domain CFMs may arise from two concurrent mechanisms: short-range interactions between *cis*-regulatory elements and loop extrusion between nearby CTCF sites. The contribution of *cis*-regulatory interactions is supported by the observation that domain CFM usage varies markedly between cell types, within and between tissues, and under metabolic perturbations that alter transcription factor levels. In contrast, our data support a predominant role of loop extrusion in the formation of short-range loops as Rad21-degradation markedly reduces domain CFMs in human cells, yet changes in TF composition and levels between cell types or during metabolic disease do not alter CFMs but rather change the frequency at which they appear in single cells. Together, these findings suggest that while both mechanisms participate in short-range loops, loop extrusion provides the dominant architectural scaffold, whereas *cis*-regulatory interactions primarily modulate motif usage within this shared structural repertoire.

Consistent with the limited processivity of loop extrusion factors, distant loops within TADs are significantly and systematically less common (5-15%) than short-range loops (∼15-25%). These measurements provide direct *in vivo* estimates of looping frequencies, extend recent live-imaging estimates of CTCF looping frequencies in cultured cells (3–30%, depending on genomic distance between CTCF anchors)^24,25^, and quantify loop frequencies for tens of loops across multiple loci, tissues and species, providing a broad and comprehensive view of extrusion-driven contacts at regulatory scales.

Most single traces contain combinations of CFMs rather than a single dominant motif. Co-occurrence between distinct loop CFMs is rare, indicating that most alleles contain at most one distal looping configuration at a time. In contrast, loop CFMs frequently co-occur with domain CFMs, suggesting that distal looping can coexist with local domain-like interactions within the same allele. Because 3DTopic captures the most prominent recurrent motifs, other conformations such as partially extruded intermediates^7,68^ are likely present but are recovered less frequently due to their structural heterogeneity and transient nature. Importantly, the restricted combinations of CFMs observed across alleles indicate that chromatin conformations explore a structured subset of possible configurations, consistent with the low-dimensional nature of the conformational space.

Third, why are enhancer-promoter distances higher in transcribing cells? The canonical view of enhancer action involves physical proximity to their target promoters^75^. However, this model was recently challenged by the observation that enhancers can be more distant from their target promoters in transcribing cells^31,76^. Our experiments reveal a similar phenomenon: in beta cells, domain and loop CFMs involving *Pdx1* and its proximal or distal enhancers occur less frequently than in non-transcribing exocrine cells. However, in beta cells *Pdx1* not only increases its distance to its enhancers but also to most loci within its TAD. Together, these observations indicate that transcriptional activation is accompanied by locus decompaction, in which *Pdx1* expands and becomes physically isolated from its surrounding chromatin. This interpretation is consistent with previous work on the formation of transcription loops by long, highly expressed genes^77^. In contrast, models invoking the formation of condensate-like micro-environments^78^ would predict frequent co-occurrence of enhancer-promoter contacts and motifs linking multiple distant regulatory elements in transcribing cells. Our data do not support these predictions, suggesting that in our system, the formation of stable multi-element regulatory assemblies are rare, and that enhancer-promoter distances primarily reflect transcription-dependent gene decompaction.

Fourth, is 3D TAD organization in single cells conserved? Our results show that multiple CFMs at the *Pdx1/PDX1* locus identified in mouse pancreas are also detected in human fetal pancreas, particularly CFMs that involve *Pdx1/PDX1* and its proximal and distal regulatory elements. This cross-species similarity indicates that single alleles in both species explore comparable conformational landscapes. Notably, the *Pdx1/PDX1* loci in mouse and human share a similar arrangement of *cis*-regulatory elements and CTCF binding sites, suggesting that this regulatory architecture is sufficient to generate similar chromatin folding motifs. Surprisingly, these observations support the idea that the repertoire of motifs is largely preconfigured by the underlying genomic organization of regulatory elements.

While our analyses reveal organizational principles across tissues and species, these conclusions are based on a limited number of genomic loci. The loci examined differ in their 1D genomic architecture, including the density and positioning of genes, CTCF sites, and *cis*-regulatory elements, which may influence their local folding behavior. Accordingly, differences in CFM usage between cell types likely reflect, at least in part, the distinct transcriptional activity and regulatory landscape of each locus. Future extensions of this approach to larger genomic panels and additional tissues will be required to determine the generality of these observations and to identify potential locus-specific versus universal folding patterns.

Finally, can disease states alter 3D TAD organization in single cells? While single-cell transcriptomic studies have revealed that many diseases originate from transcriptional dysregulation^79^, whether such changes are mirrored by alterations in single-cell 3D chromatin structure has remained unclear. Our results demonstrate that the overall ensemble of 3D structures explored by the *Pdx1* TAD displays detectable changes in a type 2 disease model. Most notably, these differences arise from selective motif reweighting in defined subpopulations of alleles, rather than uniform remodeling across all beta-cell traces. A subset of *Pdx1* CFMs becomes more prevalent in mice with type 2 diabetes, resembling the CFM changes observed in exocrine cells where *Pdx1* is poorly transcribed, and consistent with the depletion of *Pdx1*^39,63,80^. However, the precise pattern of CFM alterations differs from that of exocrine cells, showing that HFD beta cells do not simply adopt an exocrine-like folding state despite reduced *Pdx1* transcription in both contexts, and in line with evidence that obesity and T2D induce a partial dedifferentiation of beta cells into nonfunctional endocrine progenitor-like cells or promote their transdifferentiation into alternative endocrine types^66^. These analyses underscore the power of high-resolution single-allele chromatin imaging and unsupervised analyses, as conventional ensemble approaches would average out the subtle yet functionally relevant changes in 3D genome organization that may accompany a disease state.

In summary, we provide a blueprint for using imaging-based spatial genomics to chart 3D chromosome organization in single cells across multiple cell types, and apply machine learning to unveil recurrent motifs in 3D chromatin organization in different healthy tissues and during disease. These technologies will be critical to further dissect the role of 3D chromatin structure in the regulation of cell type specific transcriptional programs, and the molecular mechanisms involved.

## Methods

### Mice

Animal studies were conducted according to the European Animal Welfare Guidelines (2010/63/EU). Protocols were approved by the Institutional Animal Care and Use Committee (CEEA-LR-1434) and the French Ministry of Agriculture (APAFIS#13044). Mice were housed in a conventional facility on a 12 hour light/12 hour dark cycle and were given chow and water ad libitum. C57BL/6J mice were purchased from Janvier-SAS (RRID: IMSR_JAX:000664). Mice were fed with normal diet (ND) until 6 weeks of age and then fed with either ND or high-fat diet (HFD) (63% energy from fat) (Safe Diets, France) for 14 weeks. No data were excluded unless animals died during experimentation. The intraperitoneal glucose tolerance test (IPGTT) was performed as described^81^. We chose a glucose dose of 3 g/kg body weight to ensure full beta cell challenge and generation of an insulin peak^82^.

### Mouse and human tissue sample preparation

Mice were anesthetized with ketamine/xylazine and perfused with cold PBS through the intracardiac route. The collected tissues were then fixed in 4% paraformaldehyde in PBS (v/v) (Sigma) at room temperature for 4 hours under agitation, followed by overnight incubation at 4°C. Subsequently, the tissues were transferred to a PBS solution containing 30% sucrose until they sank to the bottom of the tube. Tissues were then embedded in OCT (Sigma) and stored at –80°C until cryosectioning. Prior to cryosectioning, 40 mm round coverslips (Bioptechs) were washed with 70% ethanol in water (v/v) and activated with air plasma for 30 seconds. Slides were then covered with 100 µL of pure 3-aminopropyltrimethoxysilane (Sigma) for 5 minutes at room temperature. Slides were left in water for 5-10 minutes and rinsed 2×10 minutes in water with agitation. A solution containing 0.5% glutaraldehyde (Sigma) in PBS was added for 30 minutes and rinsed with water. The slides were then coated with 0.1 mg/mL poly-D-lysine (Sigma) in water for 1 hour and incubated O/N in water. Ten µm tissue sections were cut with a cryostat and added to the coated slides, dried at room temperature for 1-2 hours for immediate use, or frozen at –20°C for later use.

Human fetal pancreatic tissues (post-conceptional week 9-11) were obtained through the Inserm cross-cutting scientific programme Human Developmental Cell Atlas (HuDeCA) from surgical abortions in accordance with the French bioethics legislation and Inserm guidelines as in Ref.^83^. Maternal written consent was obtained, along with approval from Agence de Biomédecine, the competent authority in France. Dissected fetal pancreatic tissues were fixed by immersion in 4% paraformaldehyde in PBS (v/v) (Sigma) at 4°C for 1 to 5 days, depending on size. Subsequently, tissues were transferred to a PBS solution containing 30% sucrose until they sank to the bottom of the tube, embedded in OCT (Sigma) and stored at –80°C until cryosectioning. Human tissues were then processed as described above for mouse tissues.

### RNA-FISH libraries

Sequential RNA-FISH libraries were constructed following the procedure described in Ref.^84^ using the library design script from https://github.com/ZhuangLab/MERFISH_analysis. Briefly, a maximum of 90 single-stranded DNA probes were designed with a homology region of 30 bp, allowing 20 bp overlap between adjacent probes to maximize the number of probes for each transcript. Each probe was customized to add a tail composed of a transcript-specific readout sequence and a forward and reverse primer sequence positioned at the 5’ and 3’ ends for PCR amplification of the library following the method described in Ref.^45^. Validation was performed by immunofluorescence staining and by verifying that the smFISH patterns of *Ins1*, *Gcg* and *Sst* staining corresponded to the expected core-mantle arrangement of endocrine markers. The full list of readout and probe sequences is available in Supplementary Data 2 and 3, respectively.

### Hi-M libraries

Oligopaint libraries were obtained from a public database (http://genetics.med.harvard.edu/oligopaints) containing unique 35/45mer sequences homologous to the Mouse genome (mm10) or to the Human genome (hg19). Hi-M probe sets were designed for each locus and amplified according to the method described by Cardozo Gizzi et al. (2020). Four mouse Hi-M libraries were designed. The first two libraries encompassed the *Pdx1* gene: one containing 25 barcodes of approximately 25 kb each, named *Pdx1* low-resolution library (Pdx1 LR), and one containing 18 barcodes of approximately 8 kb each, named *Pdx1* high-resolution library (Pdx1 HR). The third library was designed along the *Ins2* gene and contained 25 barcodes, each targeting regions of approximately 6 kb. The final library was designed along the *Isl1* gene and consisted of 24 barcodes, each targeting regions of approximately 26 kb. In addition, one human Hi-M library was designed for the *PDX1* locus, comprising 23 barcodes each covering approximately 29 kb. The full list of barcode and probes sequence is available in Supplementary Data 1 and 2.

### DNA-FISH labeling

Slides were treated with RNase A at 200 µg/ml in PBS for 1 hour at RT before incubation in Sodium Citrate (10mM citrate, 0.05% Tween, pH 6.0) for 5 minutes at RT. Next, slides were incubated with 10 mM sodium citrate for 25 minutes at 80°C in a water bath and left on the bench for 1h. Slides were washed with 2xSSC 5 minutes and incubated in 50% formamide wash buffer for 2 hours at RT. Slides were placed upside down on glass Petri dishes in contact with 2 µL of 5 to 10 µg\µL library, 1 µL of 100 µM of the fiducial library, in 20 µL of hybridization buffer (FHB) (50% formamide, 10% dextran sulfate, 2xSSC, 0.5 mg/mL Salmon Sperm DNA). Slides were incubated for 3 hours at 45°C in a water bath before a heat-shock at 85°C for 5 minutes on a heating block. Slides were then incubated in a humidity-controlled 37°C incubator overnight. Slides were then washed under agitation at 80 rpm during40 minutes in 50% formamide wash buffer twice, 20 minutes in 40% formamide wash buffer, 20 minutes in 30% formamide wash buffer, 20 minutes in 20% formamide wash buffer, 20 minutes in 10% formamide wash buffer and 20 minutes in 2xSSC. Slides were post-fixed with 4% PFA for 10 minutes at RT and stored in 2xSSC at 4°C until imaging. The same procedure was applied to both mouse and human samples.

### RNA-FISH labeling

Slides were brought to RT and left 1 hour to dry. Tissues were post-fixed for 10 minutes with 4 % PFA and washed with PBS. Slides were incubated in cold 70% EtOH overnight at 4°C. Slides were then rehydrated in 2xSSC for 5 minutes at RT and incubated for 3 hours in a wash buffer (30% formamide in 2XSSC) at 37°C. Slides were placed upside down on glass Petri dishes in contact with 2 µL of 5 to 10 µg/µL ssDNA library diluted in 20 µL of hybridization buffer (HB) (30% Formamide, 10% dextran sulfate, 2X SSC, 500 µL tRNA stock (20 mg/ml), 100 µl of RVC stock (200 mM)) and incubated in a humidity-controlled 37°C incubator for 36 hours. Slides were then washed for 30 minutes with a 30% formamide wash buffer at 45°C twice and stored in 2xSSC at 4°C until imaging.

### Imaging system

Experiments were performed on two similar acquisition setups. A home-made imaging setup built on a RAMM modular microscope system (Applied Scientific Instrumentation), as described previously^45,85^ and a modified Zeiss microscope (Axio Observer) with a similar design. Both setups were equipped with a water immersion objective (Nikon Plan-Apo 60x NA=1.2 and Zeiss W DICII 63x NA=1.2 respectively), a sCMOS camera (Flash4-v3, Hamamatsu) and a home-made fluidic system.

### Acquisition of Hi-M datasets

Slides were mounted in an FCS2® flow chamber (Bioptechs, USA). Approximately 20-30 regions of interest (ROI) were selected and imaged (215 x 215 µm – pixel size 105 nm). A mix containing an adapter complementary to the reverse primer sequence was injected for “mask0” imaging (25 nM in 2×SSC, 50% v:v formamide). Slides were washed with 2.8 mL of wash buffer (2×SSC, 50% v:v formamide) and incubated for 15 minutes with a readout hybridization mix (25 nM of Atto-488 imager probe complementary to the fiducial library, 25 nM Alexa-SS-647-coupled probe complementary to the “mask0” adapter, 2×SSC, 50% v:v formamide). Slides were then washed with 2.8 mL of wash buffer solution (2×SSC, 50% v:v formamide) and flushed with 1.5 mL of 0.5 µg/ml of DAPI solution in 2×SSC to stain nuclei. A 3-color stack of images (z-step size of 250 nm, 20 µm stack) was acquired for DAPI (405 nm), the fiducial library tagged with Atto488 (488 nm) and the “mask0” tagged with Alexa647 (640 nm). After imaging, the chambre was flushed with 1 mL of chemical bleach buffer (2×SCC, 50 mM tris(2-carboxyethyl)phosphine (TCEP) to bleach the “mask0” imager. The sample was then sequentially hybridized as follows. A solution containing the imager oligo (25 nM Alexa-SS-647 probe, 2×SSC, 50% v:v formamide) was injected and incubated for 15 minutes. Slides were then washed with 2 mL of wash buffer and with 1 mL of 2xSSC before injecting the imaging buffer (IB) (1xPBS, 5% w/v glucose, 0.5 mg/mL glucose oxidase and 0.05 mg/mL catalase). In each cycle, fiducials and readout probes were sequentially imaged with 488 nm and 647 nm excitation lasers respectively. After imaging, the fluorescent tag of the readout probes was cleaved using 1 mL of chemical bleach buffer (2× SCC, 50 mM TCEP hydrochloride). Finally, the samples were washed with 1 mL of 2×SSC before a new hybridization cycle started.

### Combined seq RNA-FISH and Hi-M

After seq RNA-FISH experiment, the slide was removed from the FCS2® flow chamber and DNA-FISH labeling protocol was performed. The slide was then mounted again in the FCS2® flow chamber and realigned with micron precision using triangulation marks on both the coverslip and the flow chamber glass. Pixel accuracy was achieved by correcting the TIFF images by cross-correlating the DAPI staining acquired during RNA and Hi-M acquisitions using a custom Python script.

### Immunostaining

Pancreas frozen slices (10 µm) were labeled with antibodies as described in Schaeffer, *et al.*^86^. Images were acquired using a Zeiss LSM 800 confocal microscope. For quantifications, three slices were randomly selected from at least two animals per group. Antibodies used were: rabbit anti-CTCF (1:150, Cell Signaling 3418), anti-rabbit Cy3 (1:1000, Jackson Immunoresearch 711-165-152). Nuclei were labeled using DAPI (Sigma). To quantify fluorescence in nuclei following CTCF labeling, nuclei were segmented in 3D using Cellpose^87^. The median fluorescence signal was computed per nucleus and the average signal was computed for each islet.

### Image analysis

TIFF images were then deconvolved using Huygens Professional 21.04 (Scientific Volume Imaging, https://svi.nl). The analysis was performed using the pyHiM analysis pipeline we developed (https://pyhim.readthedocs.io/en/latest/) with release 0.7^88^. To obtain proximity frequency maps from pairwise distance matrices, we used a distance threshold of 100 nm, a value that optimizes the correlation between Hi-M and Hi-C datasets (**Figs. S1h-i**).

### Cell type assignment to Hi-M traces

Cell type assignment was performed using RNA-FISH signals according to the following pipeline. First, for segmentation of alpha, beta and delta cells, specific RNA-FISH signals were acquired for each cell marker, *Gcg*, *Ins2* and *Sst*. These images were segmented in Ilastik^89^ to define a 3D mask for each cell type. In a second step, RNA-FISH masks and Hi-M DNA-FISH traces were registered based on the DAPI signal (see section *Combined seq RNA-FISH and Hi-M*) using a custom Python script. Finally, traces were assigned to specific cell types according to the 3D RNA-FISH mask with which they overlapped. Exocrine cells were identified as cells that were never assigned to any of the aforementioned RNA-FISH masks.

### Trace normalization and imputation

Variability in 3D distance distributions can sometimes be observed between experiments and is likely to be due to small variations in fixation conditions, Hi-M labeling and acquisition conditions. To minimize the effect of these variations on Hi-M pairwise distance matrices, the following normalization procedure was used:first, the physical vs. genomic distance curves are calculated for each experiment. By pooling all data, an average curve is also calculated. To correct for experimental variability, we computed a correction factor for each experiment by determining the scaling value that minimized the deviation between the individual curve and the global average curve. This correction factor was then used to align each experiment’s distance measurements to the global reference.

Additionally, missing barcodes in Hi-M traces were imputed as follows: If a barcode was missing in a Hi-M trace, but the two adjacent barcodes were present, the position of the missing barcode was imputed by averaging the positions of the two adjacent barcodes.

### Volcano plots

Volcano plots from Hi-M matrices were computed for two conditions. First, the differential median PWD matrix was calculated by determining the ratio of each pair of bins between the two conditions. Then, p-values were derived using the Wilcoxon rank-sum test, comparing the single-cell distance distributions for each pair of bins. Finally, the log2 values of the differential matrix were plotted against the –log10(p-values) from the Wilcoxon test.

### 3DTopic modeling

To apply 3DTopic on Hi-M data, we used the <u>LDA implementation</u> of the scikit-learn Python package. Other Python packages were tested^90^ which gave identical results.

LDA was first introduced in Natural Language Processing (NLP) to analyze large corpus of text documents and extract the most frequently discussed topics within these documents. LDA is a Bayesian model that associates words within the same topic according to their co-occurrence, i.e. the probability that these words occur together in the same text. Topic decomposition was recently used to find cis-regulatory regions from ATACseq data^91,92^. Similarly, for the Hi-M data analysis, we used LDA to decompose single cell pairwise contact maps into a discrete number of topics, which we called Chromatin Folding Motifs (CFMs). Similar to text analysis, each CFM consists of a specific set of contacts that have a high probability to be detected together in the Hi-M data.

In order to perform LDA on Hi-M data, only the traces containing at least 65% of the barcodes were selected. Each selected trace was converted into a binary pairwise contact map using a distance threshold optimized to maximize the correlation between Hi-M and Hi-C datasets (Figs S1i and S10c). In the case of the *Pdx1* low-resolution library, this optimal threshold was determined to be 100 nm (see Fig. S1i): two loci were considered to be in contact (’1’) if the 3D distance separating them was less than the 100 nm threshold, otherwise the contact value was set to zero. Each map was then linearised into a single 1D vector according to the LDA input data format requirements. As the contact maps are symmetrical along the main diagonal, only the upper half of the map without the diagonal is used. For example, for the Pdx1 (low resolution) library, each 25×25 pairwise contact map is converted into a 300-bit 1D vector. Missing data were assigned a value of ‘0’, indicating the absence of contact information.

LDA was applied to all formatted traces (minimum 1500) pooled from multiple experimental replicates. We use batch mode and between 2000-4000 iterations to obtain stable topics. The number of traces used for each training is indicated in **Table 1**. Depending on the data set, between 15 and 30 topics (CFMs) were used for the decompositions. This parameter was chosen based on the maximum coherence^48^ calculated with different numbers of topics, ranging from 5 to 50 (**Figs. S2a and S10e**).

**Table 1:** Training dataset used for 3DTopic decomposition.

| Name of the training | Source of traces | Number of traces for training | Sample used for training |
| --- | --- | --- | --- |
| *Pdx1* LR | Hi-M | 6174 (each) | Pancreas endocrine, Pancreas exocrine |
| *Pdx1* LR | Hi-M | 11609 (Brain 1661 / Liver 1424 / Lung 2853 / Lymph node 2647 / Thymus 1607 / Kidney 1417) | Brain, Liver, Lung, Lymph node, Thymus, Kidney |
| *Pdx1* LR randomized | Simulation | 6174 (each) | Pancreas endocrine, Pancreas exocrine |
| *Isl1* LR | Hi-M | 1629 (each) | Pancreas endocrine, Pancreas exocrine |
| *Ins2* HR | Hi-M | 1982 | Pancreas |
| Bintu et al. | Chromatin tracing | 11547 | HCT116 (WT) |
| Bintu et al. | Chromatin tracing | 9356 | HCT116 (RAD21 depleted) |
| Block copolymer | Polymer simulation (Domain), Fig. S9a. | 7500 | N/A |
| Loop extrusion | Polymer simulation (Loop), Fig. S9b. | 7500 | N/A |
| Block copolymer *Pdx1* LR | Polymer simulation, Fig. 3a. | 5000 | N/A |
| Loop extrusion *Pdx1* LR | Polymer simulation, Fig. 3b. | 10000 | N/A |
| Loop extrusion | Polymer simulation (Domain), Fig. S9g. | 7500 | N/A |
| *Pdx1* HR | Hi-M | 3551 (each) | Pancreas endocrine, Pancreas exocrine |

Each LDA decomposition produced the following outputs::

● The optimized CFMs (e.g., a 30×300 matrix for the *Pdx1* library).
● The decomposition of each selected single cell data in the CFMs base, describing the contribution of each CFM to the observed contact pattern. Formally, each single-cell contact map was represented as a weighted sum of CFMs:

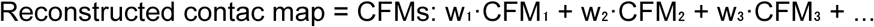

where the weights (w₁, w₂, w₃, …) reflect the contribution of each CFM to that particular map (**Fig. S4f**). For each single-cell trace, the major CFM was defined as the topic with the highest weight, representing the dominant chromatin folding motif characterizing that cell.
● A trained model that can be used to project new single-cell Hi-M datasets onto the same CFM basis.

For each dataset, multiple independent LDA runs (>10) were performed after random shuffling of the input traces to evaluate decomposition reproducibility. Finally, the best decomposition was manually selected based on its similarity to the consensus, quantified by Pearson correlation between each component’s CFM and the corresponding consensus CFM.

### Coherence computation

The coherence score is computed following the method introduced in Ref.^48^. For each topic, the Normalized Pointwise Mutual Information (NPMI) was computed among the *N* most representative contacts (typically *N = 10*, corresponding to the contacts with the highest intensity values). As detailed below, NPMI quantifies the statistical association between two contacts, Ci and Cj, by comparing the probability of co-occurrence p(Ci, Cj) with their probability assuming independence:

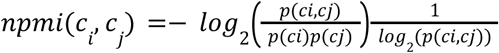

For each top contact Cn, NPMI is computed against all other top contacts in the same topic:

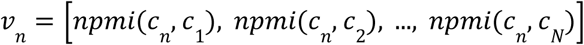

Finally, for the most relevant contacts, coherence of each topic is computed based on the average NPMI:

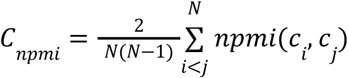

Since the number of representative contacts may vary across datasets, the range of *N*-top-contacts values was adjusted accordingly. For each dataset, an average coherence curve was computed. For instance, in the case of the *pdx1* pancreas dataset, the number of top contacts considered ranged between 5 and 10. The optimal number of CFMs used for training was then defined as the value corresponding to the maximum or plateau of the coherence curves for both methods. When the two estimates slightly differed, their average value was selected.

### UMAP based CFMs decomposition

Single-allele Hi-M traces were embedded into a low-dimensional space using Uniform Manifold Approximation and Projection (UMAP)^93^ implemented in the umap-learn Python package. Each trace was represented by its CFM decomposition vector (see section 3DTopic modeling), i.e., the set of weights describing the contribution of each Chromatin Folding Motif (CFM) to that single-allele contact map. UMAP was then applied to these CFM-based feature vectors to visualize and explore the heterogeneity of chromatin folding configurations across individual alleles. The embedding was computed in two dimensions using the following parameters, optimized to enhance data dispersion and preserve local relationships: metric=’cosine’, n_neighbors=50, min_dist=0.3, and n_components=2.

### Cell-type comparison using UMAP

To compare chromatin conformations between two cell-type conditions (e.g., exocrine versus endocrine cells), the CFMs-derived decompositions for all single cells were embedded into a shared low-dimensional space using UMAP. UMAP embeddings were computed with the following parameters: metric = ‘cosine’, min_dist = 0.3, n_neighbors = 50, n_components = 2, and random_state = 25.

We first computed a coarse two-dimensional histogram of UMAP coordinates for each condition. To estimate variability, we performed bootstrap resampling by repeatedly computing the histograms on random 50% subsamples of the data. This procedure was repeated over multiple shifted grid origins in both x and y directions to increase spatial resolution while maintaining sufficient counts per bin. For each grid shift, bootstrapped log-ratio maps were interpolated onto a common fine grid and averaged. Statistical filtering was then applied: bins for which the absolute mean log-ratio did not exceed its bootstrapped standard deviation were removed. The resulting high-resolution log10 density-ratio map highlights regions enriched (values > 0) or depleted (values < 0) for each condition.

These filtered density-ratio maps were subsequently used to identify condition-specific regions in UMAP space. For each condition, connected-component clustering was applied to contiguous enriched bins, defining distinct UMAP clusters. For each cluster, all associated single-allele contact maps were aggregated to compute a mean contact matrix, providing representative contact structures for each condition-enriched region. Importantly, all major features of the analysis were robust across a range of UMAP parameters, including changes in *min_dist* and the random initialization seed.

### Comparison of CFMs trainings

200 independent trainings were performed under identical conditions using the *Pdx1* pancreas dataset, yielding a total of 200 × 30 = 6000 CFMs. These CFMs were subsequently projected onto the reference model to obtain their respective decompositions within that framework. The rationale behind this approach is that, if a newly generated CFM closely matches one of the reference decompositions, its projection onto the reference model should yield a weight distribution characterized by a dominant component (∼1) for the corresponding topic and near-zero weights for the others. This strategy enables the assessment of topic reproducibility across trainings by embedding their reference-based decompositions into a UMAP space (metric = ‘cosine’, min_dist = 0.3, n_components = 2, n_neighbors = 50). CFMs with similar decompositions cluster together, allowing the identification and analysis of groups of CFMs that share comparable CFMs structures.

Clustering was performed in two steps: first, the HDBSCAN algorithm (method = ‘eom’, min_cluster_size = 75, min_samples = 1) was applied to identify well-defined and dense clusters. Then, a semi-supervised *LabelSpreading* algorithm was used to reassign un-assigned CFMs to the nearest clusters. CFMs with assignment probabilities below 0.8 were retained as noise to preserve clustering robustness.

### Comparison of CFMs between experimental conditions

To capture the changes induced by the perturbation of chromatin structure, chromatin tracing data Ref.^18^ in wild-type and RAD21-depleted conditions were used. 3Dtopic was first used to decompose the data into 20 and 15 CFMs for the wild-type and RAD21-depleted conditions respectively. The number of CFMs selected was based on the NPMI measure as described in the Methods section “Coherence Computation”. One hundred topic models were trained for each condition, yielding a total of 2000 and 1500 CFMs respectively after pooling. All CFMs were embedded in a two-dimensional space using a UMAP with the following manually-optimized parameters: min_dist = 0.8, num_neighbors = 12. The embedded CFMs were then colored by their source condition and maximum pseudocount. Clustering of the embedded CFMs from the wild-type condition was performed using HDBSCAN with the following optimized parameters: min_cluster_size = 40, min_samples = 2, cluster_selection_epsilon = 0.45, cluster_selection_method = “leaf”. Embedded CFMs from the RAD21-depleted condition falling within these clusters were identified by drawing a convex hull around the wild-type clusters using scipy.spatial.ConvexHull. Consensus CFMs were computed for each cluster by taking the average of embedded CFMS within and visualized for comparison between the two conditions.

### 3Dtopic reconstructions

Each single-allele decomposed matrix was reconstructed by multiplying the weight of each CFM by its corresponding CFM matrix. The reconstructed single-allele matrix was then generated by summing all weighted CFM matrices for that cell. Finally, the mean 3DTopic reconstructed map was obtained by averaging the reconstructed single-allele matrices.

### Odds ratio

The Odds Ratio (OR) compares the likelihood of an event occurring in one group relative to another. OR is calculated using the following formula:

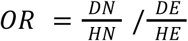

where *DN* and *DE* are the number of cells for which a given CFMs weight exceeds a minimum threshold (0.2) in test and reference tissue respectively. *HN* and *HE* are the number of cells for which a given CFMs weight does not exceed the minimum threshold (0.2) in test and reference tissue respectively.

To generate the statistical tests and the volcano plots of OR (**Figs. 5f, S14h** and **S14j**), we performed over 100 independent 3DTopic decompositions. CFMs from these decompositions were then clustered based on similarity to the reference decomposition, retaining only those with a Pearson correlation greater than 0.8. For each “similar” CFM in each decomposition, we calculated the odds ratio in the test relative to the reference tissue, yielding a distribution of the OR values for that CFM across decomposition. This OR distribution was directly displayed (**Fig. S13e**) or used to generate the volcano plots (**Figs. 5f, S14h** and **S14j**). Specifically we computed the –log10 of the p-values from a Wilcoxon test applied to the OR distribution (y-axis) and the log10 of the median OR value (x-axis).

### Consensus CFMs

To test the robustness and reproducibility of the 3D topic decomposition, consensus CFMs were computed. Multiple independent decompositions (n>10) are pooled and similar CFMs (with a Pearson correlation > 0.62) are clustered together. If a cluster contains at least n/2 topics, i.e. these topics are found in at least half of the decompositions, a consensus CFMs is derived by calculating the mean of the individual CFMs grouped within the selected cluster.

### Domain and loop CFM classification

CFMs were classified into two categories: domains and loops, using a custom Python script. Briefly, a binary mask is created based on thresholding. Connected components within the mask are analyzed for their shape and size. CFMs are categorized based on the presence and characteristics of these components: domains are identified by diagonal components, and loops by compact non-diagonal components. When necessary, visual examinations are applied to correct for misclassifications.

### CFM anchor-enrichment analysis

After training an LDA model using a dataset, all CFMs (or their average representations) obtained were linearized forms of the upper triangle matrix of the data. The CFMs were extracted and expanded into full N x N symmetric matrices using Scipy’s spatial.distance.squareform function. High-confidence contacts within a CFM were defined as the pixels whose values fell in the top 3% of the intensity distribution (bright-pixel *mask*, α = 0.97); this percentile threshold is scale-invariant and was kept fixed for all analyses.

To relate contact patterns to CTCF binding, we used a JSON annotation that listed the genomic bins occupied by CTCF from experimental data. Every bright pixel (i,j) was classified into one of two edge types: **CTCF–CTCF** or **other**. For each CFM, we built a 2 × 2 contingency table comparing the number of bright versus non-bright pixels that belonged to a given edge type κ versus all other types, and computed the Fisher exact odds ratio

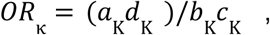

where *a*_Κ_ is the count of bright κ-edges, *b*_Κ_ bright non-κ edges, *c*_Κ_ dim κ-edges and *d*_Κ_ dim non-κ edges. Odds ratios > 1 therefore indicated enrichment of CTCF–CTCF among the strongest contacts of the CFM. Only CTCF–CTCF edge types with *OR*_Κ_ > 1 are reported on the figures presenting the analysis on Bintu *et al.*, 2018 data.

All steps were implemented in Python 3.13.5 with NumPy 2.2, SciPy 1.16 and scikit-learn 1.7.

### Single-allele trace decompositions using pre-trained models

For each genomic locus (*Pdx1* or *Ins2*), an LDA model was trained on the pancreas Hi-M dataset to define a reference set of Chromatin Folding Motifs (CFMs). The trained model was subsequently used to decompose new single-cell Hi-M datasets from the same locus but originating from different tissues (e.g., brain or lung) into the established reference CFM basis. For each input trace, the model outputs a set of weights describing the contribution of each CFM, effectively representing the contact map as a weighted linear combination of reference motifs. This transfer approach enables consistent comparison of chromatin folding configurations across datasets within a common latent space. Prior to decomposition, only traces containing at least 50% of the expected barcodes were retained.

To compare consensus CFMs between mouse and human at the Pdx1 locus, we developed a computational framework to project mouse CFMs onto the human genome. Consensus CFMs were first computed independently from mouse and human Hi-M datasets. To align mouse genomic regions (barcodes) with their human counterparts, we used the UCSC LiftOver tool to convert mouse genome coordinates (mm10) to human coordinates (hg19). For each mouse barcode present in the Hi-M library, corresponding human barcodes were identified by comparing genomic intervals. Each matrix entry (i,j) in the mouse CFM was mapped to all corresponding human bin pairs (i′,j′) based on converted coordinates. The value at (i,j) was assigned to all corresponding human bin pairs (i′,j′) when multiple mouse entries mapped to the same human pair, values were averaged. The resulting matrix represents the inferred human CFM, which approximates the expected mouse CFM projected onto the human genome. To assess CFM conservation between species, we computed the Pearson correlation between each inferred human CFM and the observed human CFM.

### Hi-C, ChIP-seq, ATAC-seq, CUT&Tag, RNA-seq and scRNA-seq data processing

ATAC-seq data were reanalyzed from raw data^94,95^. Sequencing reads were aligned to the reference mus musculus genome assembly (mm10) using Burrows-Wheeler Aligner (0.7.17-r1188) with default parameters. Finally, peak calling was performed using MACS2 (2.2.7.1) with default parameters. CTCF ChIP-seq were reanalyzed from raw data^95^ with default parameters. CTCF CUT&Tag data were downloaded from https://data.mendeley.com/datasets/mwgxv7m927/2^96^ in. bw and converted to .bedgraph using bigWigToBedGraphNext. Peaks were then called using *macs2 bdgpeakcall* with –c option set to 3. Hi-C data acquired in different mouse tissues were downloaded in .mcool format from accession number GSE150704^77^.

Annotated single-cell RNA sequencing (scRNA-seq) data were downloaded from the Tabula Muris GitHub repository (https://github.com/czbiohub-sf/tabula-muris)^97^. These data were re-plotted using a custom Python script. Quickly, the annotated data were filtered to select only the tissues used in this study, specifically Brain, Kidney, Lung, Liver, Lymph node, Pancreas, and Thymus. Within the brain tissue, both Brain_Non-Myeloid and Brain_Myeloid categories were combined and referred to as Brain. Next, tSNE1 and tSNE2 coordinates for cells from these tissues were used to represent the data in a 2D space. Expression count data were downloaded from Figshare (https://ndownloader.figshare.com/articles/5829687/versions/4) and used to represent gene expression levels in single cells within the tSNE space.

The relative expression of genes in exocrine and beta cells was determined by calculating the ratio between the count of the target gene, and the count of a housekeeping gene: Actb, in each single cell. Cells were classified as exocrine or beta cells based on the “cell_ontology_class” in the annotated scRNA-seq data. Type B pancreatic cells are referred to as beta cells, while pancreatic acinar cells are referred to as exocrine cells.

### Polymer simulations

All polymer simulations were performed using the OpenMM-based polychrom package^7,98,99^ developed by the Open Chromosome Collective (open2c, https://doi.org/10.5281/zenodo.3579473). As in Fudenberg et al.^7^, the chromatin segment corresponding to the locus was modeled as a bead-spring chain with some stiffness and excluded volume, where each bead (or monomer) was considered to be 1 kb.

For block copolymer simulations, two types of monomers were used: those corresponding to accessible sites from ATAC-seq data were assigned *type 1*, and all the others, *type 0*. We then used a smoothed square well potential (equations further below, parameters in **Table 2**) to introduce a distance-dependent attraction between all non-bonded *type 1* monomers (i.e., attraction between all accessible sites).

**Table 2:** Chromatin accessibility from ATAC-seq data and corresponding monomer in the heteropolymer chain.

| Genomic coordinates (mm10) | Monomer index | Peak intensity |
| --- | --- | --- |
| chr5:146904628 | 33 | 13.32675 |
| chr5:146912465 | 41 | 10.64644 |
| chr5:146914562 | 43 | 15.86417 |
| chr5:146938315 | 66 | 24.8245 |
| chr5:146948941 | 77 | 37.07606 |
| chr5:146962724 | 91 | 5.16083 |
| chr5:146963723 | 92 | 21.9748 |
| chr5:146974654 | 103 | 5.22891 |
| chr5:146974871 | 103 | 16.23728 |
| chr5:146983414 | 111 | 19.55906 |
| chr5:146989269 | 117 | 37.2411 |
| chr5:147017700 | 146 | 50.36518 |
| chr5:147033647 | 162 | 40.74204 |
| chr5:147043871 | 172 | 3.46557 |
| chr5:147051421 | 179 | 6.82104 |
| chr5:147055091 | 183 | 26.61741 |
| chr5:147055923 | 184 | 9.32849 |
| chr5:147057905 | 186 | 22.96821 |
| chr5:147059576 | 188 | 7.18537 |
| chr5:147064125 | 192 | 4.71837 |
| chr5:147073985 | 202 | 10.45092 |
| chr5:147075830 | 204 | 9.00849 |
| chr5:147076618 | 205 | 42.50082 |
| chr5:147124713 | 253 | 14.04529 |
| chr5:147188735 | 317 | 7.13072 |
| chr5:147227544 | 356 | 21.44765 |
| chr5:147236120 | 364 | 10.3808 |
| chr5:147258447 | 387 | 18.14113 |
| chr5:147258812 | 387 | 32.76999 |
| chr5:147262293 | 390 | 5.23312 |
| chr5:147268377 | 396 | 5.16464 |
| chr5:147270113 | 398 | 37.84033 |
| chr5:147283308 | 411 | 36.66473 |
| chr5:147293366 | 421 | 31.09091 |
| chr5:147294314 | 422 | 22.86171 |
| chr5:147299353 | 427 | 26.68709 |
| chr5:147319841 | 448 | 8.72425 |
| chr5:147402731 | 531 | 8.18355 |
| chr5:147408892 | 537 | 9.26759 |
| chr5:147430122 | 558 | 34.05257 |
| chr5:147430484 | 559 | 16.26295 |
| chr5:147431123 | 559 | 6.89618 |
| chr5:147431434 | 559 | 11.96539 |
| chr5:147460048 | 588 | 67.33253 |

For the loop extrusion simulations, the dynamics of cohesins/loop extruding factors (LEFs) were first simulated on a one-dimensional lattice with CTCF sites and their LEF capture rates (**Table 3, Fig. S9d**) assigned based on the peak sizes of CTCF from CUT&Tag data^96^. LEF positions were recorded over time and were used to rapidly construct simple contact frequency maps based on loops created during the trajectory, which then facilitated manual optimization of loop extrusion parameters (**Table 4**). Next, the recorded LEF positions were used to mimic loop extrusion on the homopolymer bead-spring chain described above under Langevin dynamics^100^ using molecular dynamics simulations (parameters in **Table 5**).

**Table 3:**
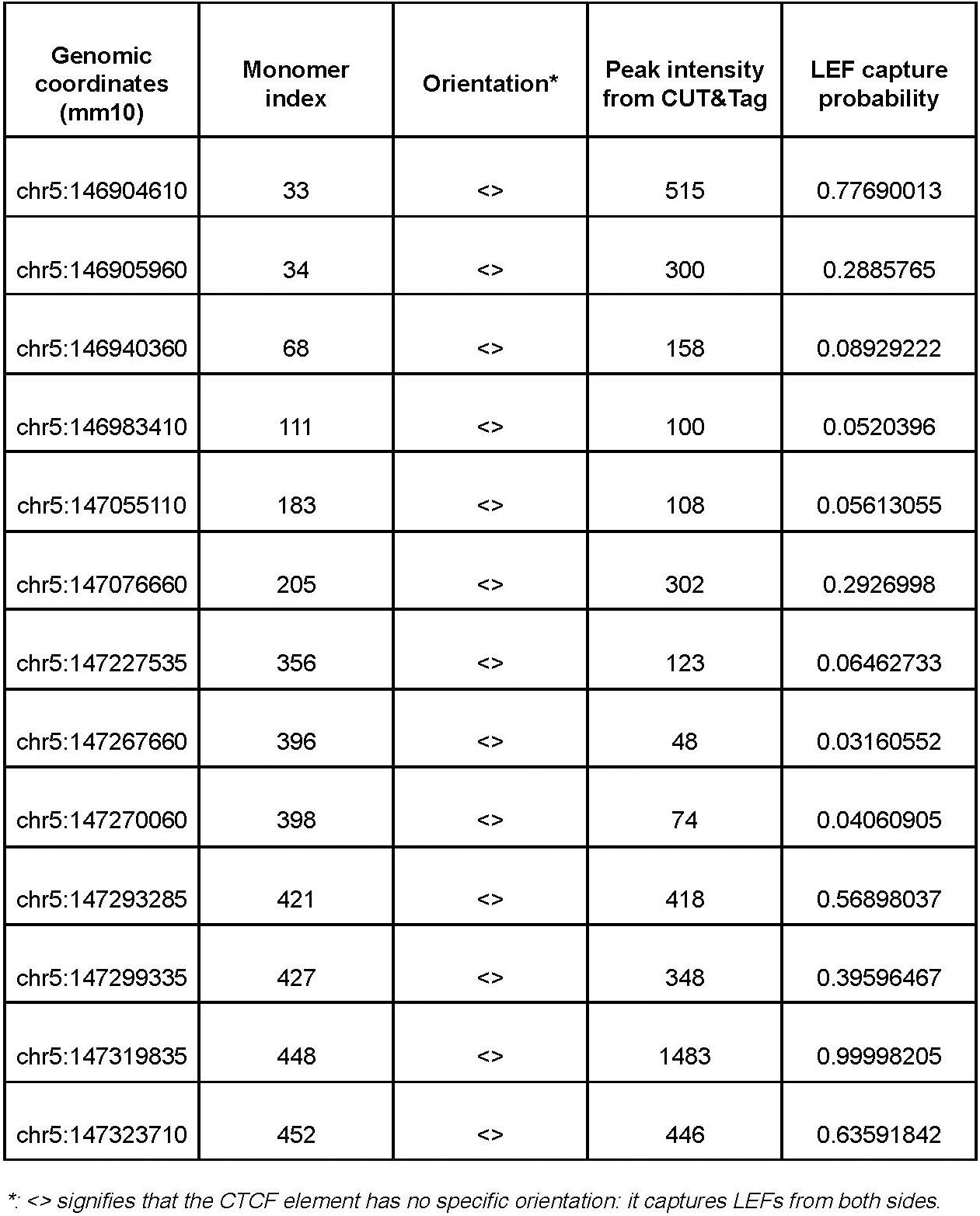
CTCF positions, orientations and their LEF capture probabilities for loop extrusion simulations.

| **Genomic coordinates (mm10)** | **Monomer index** | **Orientation\*** | **Peak intensity from CUT&Tag** | **LEF capture probability** |
| --- | --- | --- | --- | --- |
| chr5:146904610 | 33 | <> | 515 | 0.77690013 |
| chr5:146905960 | 34 | <> | 300 | 0.2885765 |
| chr5:146940360 | 68 | <> | 158 | 0.08929222 |
| chr5:146983410 | 111 | <> | 100 | 0.0520396 |
| chr5:147055110 | 183 | <> | 108 | 0.05613055 |
| chr5:147076660 | 205 | <> | 302 | 0.2926998 |
| chr5:147227535 | 356 | <> | 123 | 0.06462733 |
| chr5:147267660 | 396 | <> | 48 | 0.03160552 |
| chr5:147270060 | 398 | <> | 74 | 0.04060905 |
| chr5:147293285 | 421 | <> | 418 | 0.56898037 |
| chr5:147299335 | 427 | <> | 348 | 0.39596467 |
| chr5:147319835 | 448 | <> | 1483 | 0.99998205 |
| chr5:147323710 | 452 | <> | 446 | 0.63591842 |

\*: <> signifies that the CTCF element has no specific orientation: it captures LEFs from both sides.

**Table 4:**
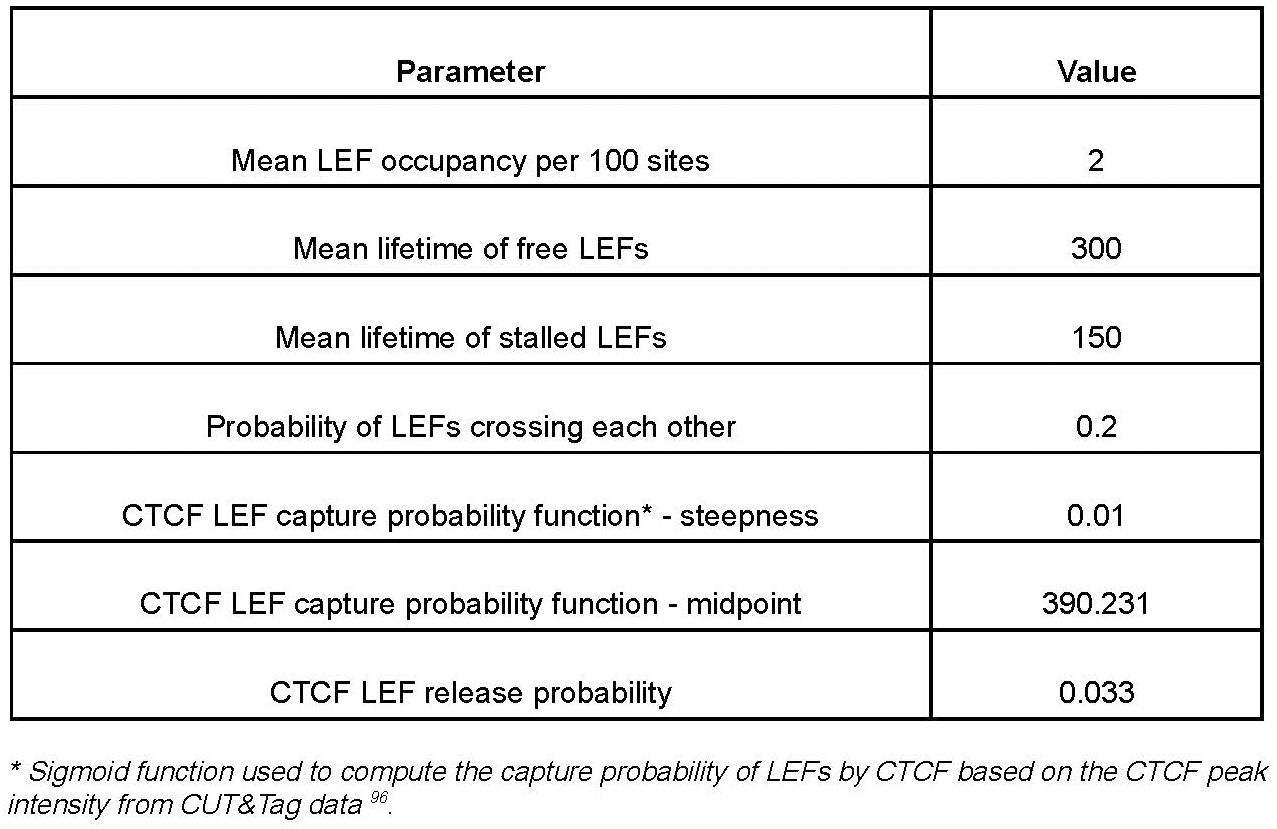
Lattice-based loop extrusion simulation parameters.

**Table 5:**
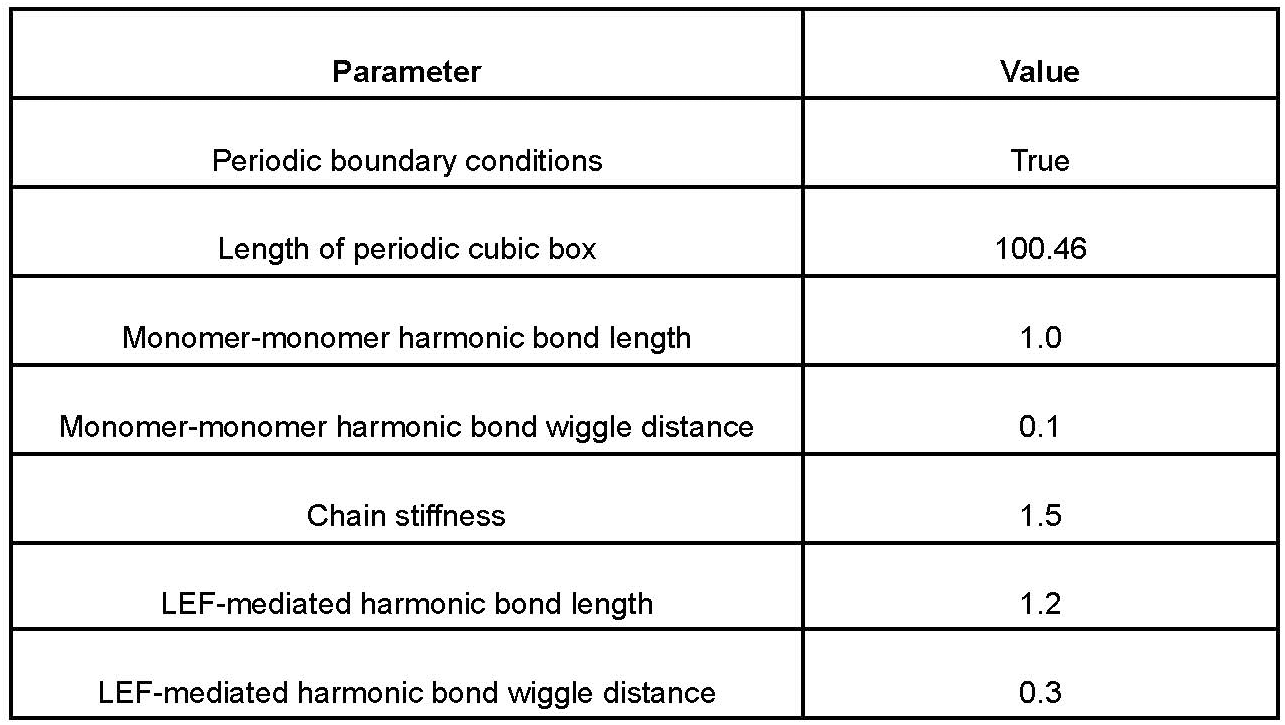
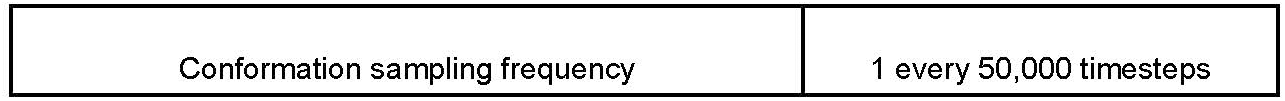
Molecular dynamics simulation parameters.

To minimize overfitting in our loop extrusion simulations, we constrained parameter choices using experimental data and prior literature. Specifically, we manually adjusted only two parameters—the number and lifetime of cohesins on the chromatin fiber—based on 1D simulations, with the sole aim of matching ensemble-averaged proximity frequency maps. The chosen values are consistent with those reported in previous studies^101^. Notably, these parameters were set *prior* to CFM detection using 3DTopic and were not optimized to reproduce experimental CFMs.

Equilibrated polymer conformations from these simulations were saved, coarse-grained to match the experimental resolution (**Table 6**), and used to construct pairwise distance matrices.

**Table 6:** Barcode target bins for polymer coarse-graining.

| Barcode ID | Target start |  | Target end |  |
| --- | --- | --- | --- | --- |
|  | Genomic coordinates (mm10) | Monomer index | Genomic coordinates (mm10) | Monomer index |
| 1 | chr5:146871445 | 0 | chr5:146898314 | 25 |
| 2 | chr5:146898349 | 26 | chr5:146921585 | 49 |
| 3 | chr5:146921627 | 50 | chr5:146943507 | 71 |
| 7 | chr5:146943542 | 72 | chr5:146961211 | 88 |
| 8 | chr5:146961252 | 89 | chr5:146983318 | 110 |
| 9 | chr5:146983361 | 111 | chr5:147003635 | 131 |
| 10 | chr5:147003670 | 132 | chr5:147023664 | 151 |
| 11 | chr5:147023746 | 152 | chr5:147041833 | 169 |
| 12 | chr5:147041970 | 170 | chr5:147061547 | 189 |
| 13 | chr5:147061586 | 190 | chr5:147083863 | 211 |
| 14 | chr5:147083907 | 212 | chr5:147109230 | 236 |
| 15 | chr5:147109291 | 237 | chr5:147136001 | 263 |
| 16 | chr5:147136044 | 264 | chr5:147170385 | 297 |
| 17 | chr5:147170433 | 298 | chr5:147195791 | 323 |
| 18 | chr5:147195829 | 324 | chr5:147223598 | 351 |
| 19 | chr5:147223719 | 352 | chr5:147244854 | 372 |
| 20 | chr5:147244895 | 373 | chr5:147270377 | 397 |
| 21 | chr5:147270412 | 398 | chr5:147293491 | 421 |
| 23 | chr5:147294086 | 422 | chr5:147310724 | 438 |
| 24 | chr5:147310794 | 439 | chr5:147331150 | 458 |
| 25 | chr5:147331233 | 459 | chr5:147356458 | 484 |
| 26 | chr5:147356493 | 485 | chr5:147378050 | 505 |
| 27 | chr5:147378091 | 506 | chr5:147414475 | 542 |
| 28 | chr5:147414513 | 543 | chr5:147450660 | 578 |
| 639 | chr5:147450695 | 579 | chr5:147499625 | 627 |

#### Potentials

All the potentials used in our simulations describe the attractive or repulsive energy between non-bonded monomers based on the distance from the center of the monomer, denoted by *r* and given in monomer-monomer bond length units.

*Homopolymer polynomial repulsive potential* (**Fig. S9b, right**)^7,98,99^

The repulsive potential used for excluded volume is given by

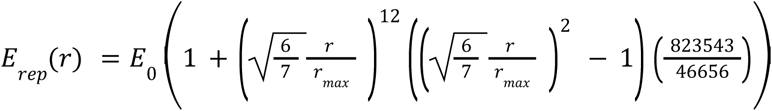

where *r*_*max*_ = 1. 05 and *E*_0_ = 3 *kT* are the width and the height of the potential respectively *Heteropolymer smoothed-square well potential* (**Fig. S9b, left**)^7,98,99^,

The repulsive part of the heteropolymer potential is given by

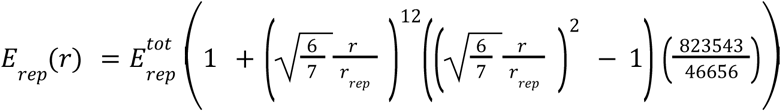

where the width of the potential *r_rep_* = 1. 05, and the total repulsive energy is

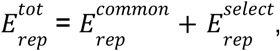

Here, 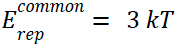 and 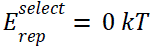 are the repulsive energies between all and selective non-bonded monomers respectively. The former accounts for excluded volume. The attractive part of the potential is described by

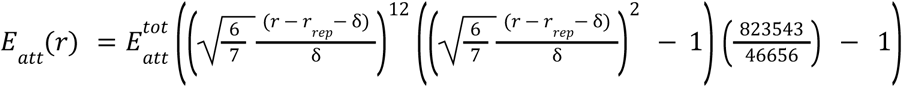

where 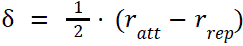, the width of the potential measured from *r* = 0 is *r_att_* = 1. 8, and the total attractive energy is

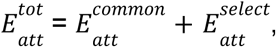

with 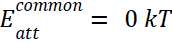 and selective attraction between *type 1* monomers 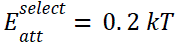

These two parts are then stitched together using a Heaviside step function

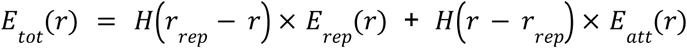

where *H*(*x*) = 0 *if x* < 0 *and H*(*x*) = 1 *if x* ≥ 0

### Comparison of experimental and simulated CFMs

Experimental and simulated CFMs were compared using Pearson correlation. In practice, each CFM map was linearized into a 1D vector, and pairwise correlations were calculated between experimental and simulated maps. For visualization purposes, simulated CFMs were then sorted from higher to lower correlation with the ranked experimental CFMs (**Fig. 3c**).

## Acknowledgements

We thank the laboratory members for the helpful discussion and feedback. We are grateful to Marcelo Rubinstein, Maria Cristina Gambetta, and David Lleres for their critical reading of the manuscript, to Alessandro Barducci for advice setting up polymer simulations, and to Julian Mozziconacci for advice related to data analysis. This project was funded by the European Union’s Horizon 2020 Research and Innovation Program (Grant ID 724429) (M.N.) and the French National Research Agency (ANR-23-CE12-0023-01) (M.N.). We also acknowledge the Bettencourt-Schueller Foundation for their prize ‘Coup d’élan pour la recherche Française’. The CBS is a member of the France-BioImaging, a national infrastructure supported by the French National Research Agency (ANR-10-INBS-04-01). O.M. was supported by an FRM and Ligue Contre la Cancer PhD fellowships. D.J.H. was supported by MRC (MR/S025618/1), Diabetes UK (17/0005681 and 22/0006389) and UKRI ERC Frontier Research Guarantee (EP/X026833/1) Grants. This work was granted access to the HPC resources of CINES under the allocation A0190316693 made by GENCI. This work was supported on behalf of the “Steve Morgan Foundation Type 1 Diabetes Grand Challenge” by Diabetes UK and SMF (grant number 23/0006627 to D.J.H. and I.A.).

## Author Contributions

O.M., M.S. and M.N. conceived the study and the design. O.M., M.S., C.E.Y acquired the data. O.M., J-B.F., M.S., C.E.Y, G.G., M.N. analyzed the data. O.M., J-B.F., G.G. and M.N. wrote analysis software. G.G and J-B.F performed loop extrusion simulations. J-B.F. built the microscope. O.M., M.S., J-B.F, G.G. and M.N. interpreted the data. M.S. and Y.K. prepared and characterized mice. I.A. and D.H. participated in experimental design. J.F. acquired and analyzed Chip-Seq datasets. J.V and R.S. provided human tissues. A.M and O.M. designed and tested RNA-FISH libraries. M.N., O.M., M.S. and J-B.F. wrote the manuscript. M.N. and M.S. supervised the study and acquired funds.

