## Supplementary Figures for "Single-allele chromatin folding is organized by recurrent motifs that are reweighted across cell types and states"

### Supplementary Information for

### “3DTopic reveals a compact basis of motifs that quantify single-allele chromatin folding across tissues, transcriptional conditions and disease”

**Table of contents**

Supplementary Data 1
Supplementary Tables 1
Supplementary Figure 1 2
Supplementary Figure 2 4
Supplementary Figure 3 6
Supplementary Figure 4 7
Supplementary Figure 5 8
Supplementary Figure 6 10
Supplementary Figure 7 12
Supplementary Figure 8 14
Supplementary Figure 9 15
Supplementary Figure 10 17
Supplementary Figure 11 19
Supplementary Figure 12 20
Supplementary Figure 13 21
Supplementary Figure 14 23
Supplementary Figure 15 25
Supplementary Figure 16 27

### Supplementary Data

SUPPLEMENTARY DATA 1. List of sequences of primary Hi-M probes.

SUPPLEMENTARY DATA 2. List of sequence of imaging (io), adapter oligos and barcodes used in this study.

SUPPLEMENTARY DATA 3. List of sequences of primary RNA-FISH probes.

### Supplementary Tables

SUPPLEMENTARY TABLE 1. List of publicly available data used in this study.

SUPPLEMENTARY TABLE 2. List of primers for library amplification used in this study.

### Supplementary Figures

**Supplementary Figure 1**

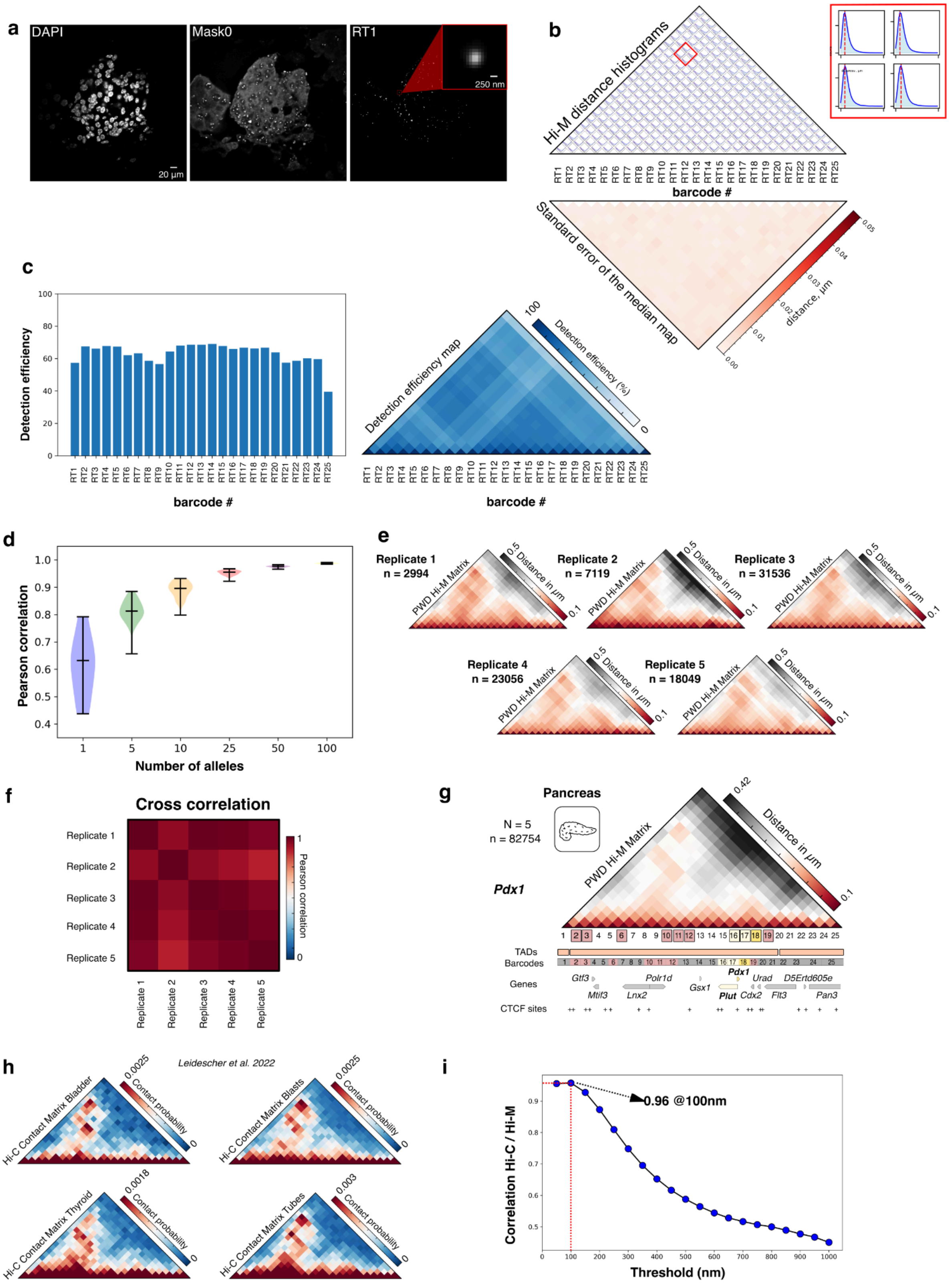


**a** Left: Grayscale DAPI-stained nuclei image of a mouse pancreas, centered on a pancreatic islet. Middle: Maximum intensity projection of the fluorescence signal from the entire *Pdx1* Hi-M library (Mask0). Right: Maximum intensity projection of the fluorescence signal from a single barcode (RT1) in the same field of view.

**b** Top: Matrix displaying histograms of pairwise distance distributions within all barcode pairs acquired on mouse pancreatic tissue. A zoom view of four histograms, highlighted in red, is shown on the right. Bottom: Matrix representing the standard error of the median for each barcode pair. Dark red and white represent high and low error respectively.

**c** Left: Bar-plot representing the detection efficiency for each barcode. Right: Detection efficiency matrix. Dark blue and white represent high and low detection efficiencies, respectively.

**d** Violin plots representing the Pearson correlation between the ensemble PWD matrix obtained using the full dataset and that using subsets containing different numbers of single allele data. Distributions were obtained by bootstrapping using 50 cycles per condition.

**e** Median PWD matrices reconstructed for the *Pdx1* locus for each individual Hi-M experiment in mouse pancreas. n indicates the number of traces used for reconstructing each map. Two animals were used.

**f** Matrix representing the Pearson correlation between each pair of Hi-M replicates in mouse pancreas.

**g** Median Hi-M pairwise distance (PWD) matrix along the *Pdx1* locus (chr5:146871445-147499625) in mouse pancreas (mm10) constructed from $n$ = 82754 traces ($N$ = 5 experiments, 2 different mice). Barcodes used for Hi-M sequential imaging are represented as color-coded boxes (pink: presence of CREs, white: *Plut* gene, yellow: *Pdx1* gene).

**h** Hi-C contact matrices from four different mouse tissues. Red and blue represent high and low contact frequencies, respectively. Data downloaded from accession number GSE150704 (*77*).

**i** Pearson correlation coefficient between the interpolated Hi-C contact map and Hi-M proximity frequency matrix was calculated for various cutoff distances. The highest correlation was achieved at a threshold of 100 nm, as indicated.

### Supplementary Figure 2

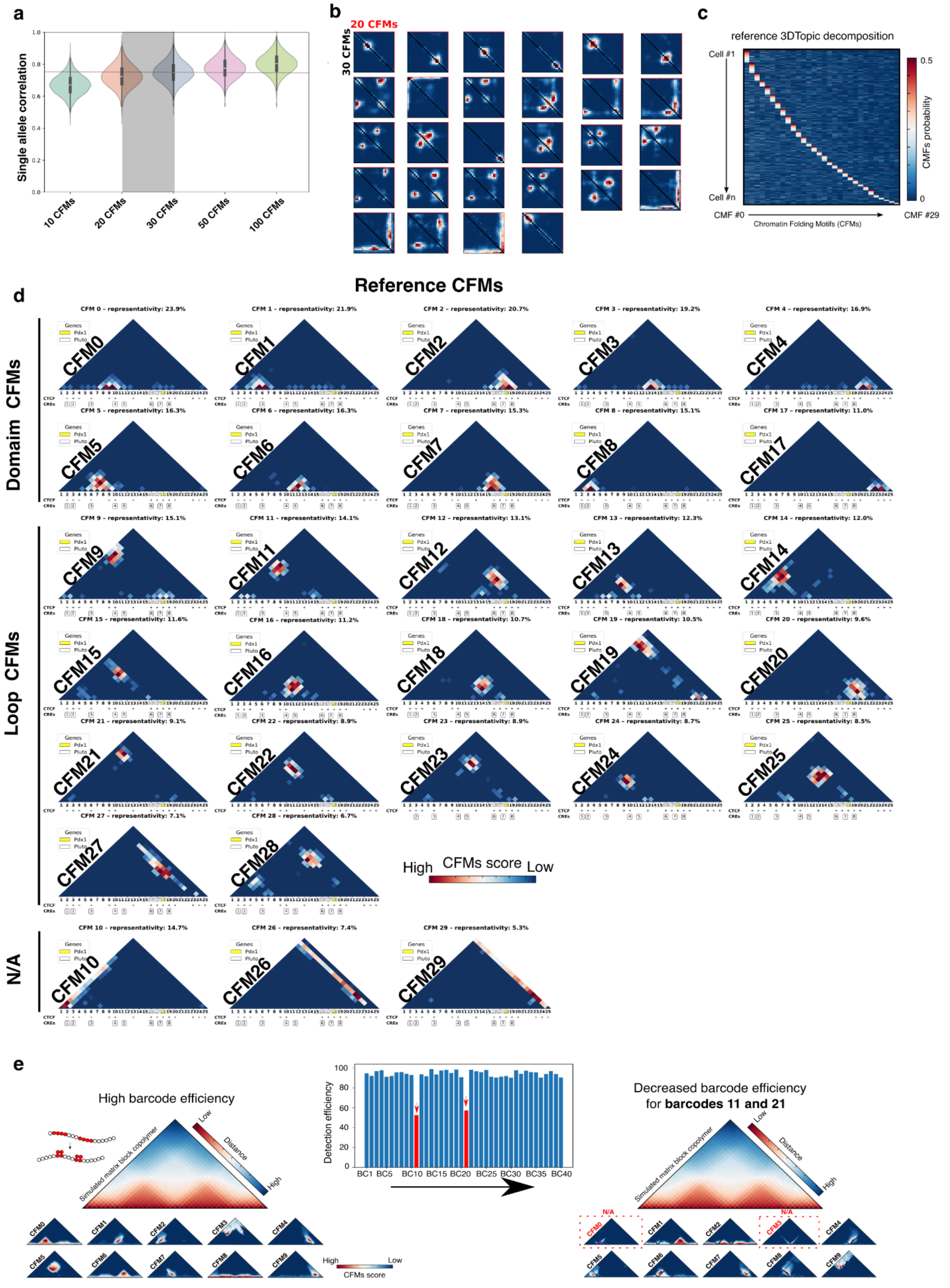


**a** Distribution of the coherence score for ten separate decompositions with varying numbers of CFMs (10, 20, 30, 50 and 100). The highest mean of coherence score was achieved with

30 CFMs, as highlighted (see *Methods*).

**b** Comparison of average CFMs per cluster for training performed with 30 CFMs (bottom-left) and 20 CFMs (top-right).

**c** Probability distribution illustrating the decomposition of pancreatic cells into 30 CFMs along the *Pdx1* locus for the reference decomposition.

**d** Complete gallery of CFMs obtained from a decomposition along the *Pdx1* locus with a cut off threshold of 100 nm. The 30 CFMs are grouped into 2 distinct categories: Domains, Loops. Structures that could not be classified into either category are labeled N/A.

**e** Left: Median PWD matrix from simulated block copolymer models with high barcode detection efficiency across single polymer trajectories, along with the derived CMFs. Middle: Bar plot showing barcode detection efficiency. Detection efficiency for barcodes 11 and 21 (highlighted in red) were deliberately reduced. Right: Median PWD matrix from simulations incorporating the reduced efficiency for Barcodes 11 and 21. The resulting CFMs display the emergence of stripe-like structures, classified as N/A.

**Supplementary Figure 3**

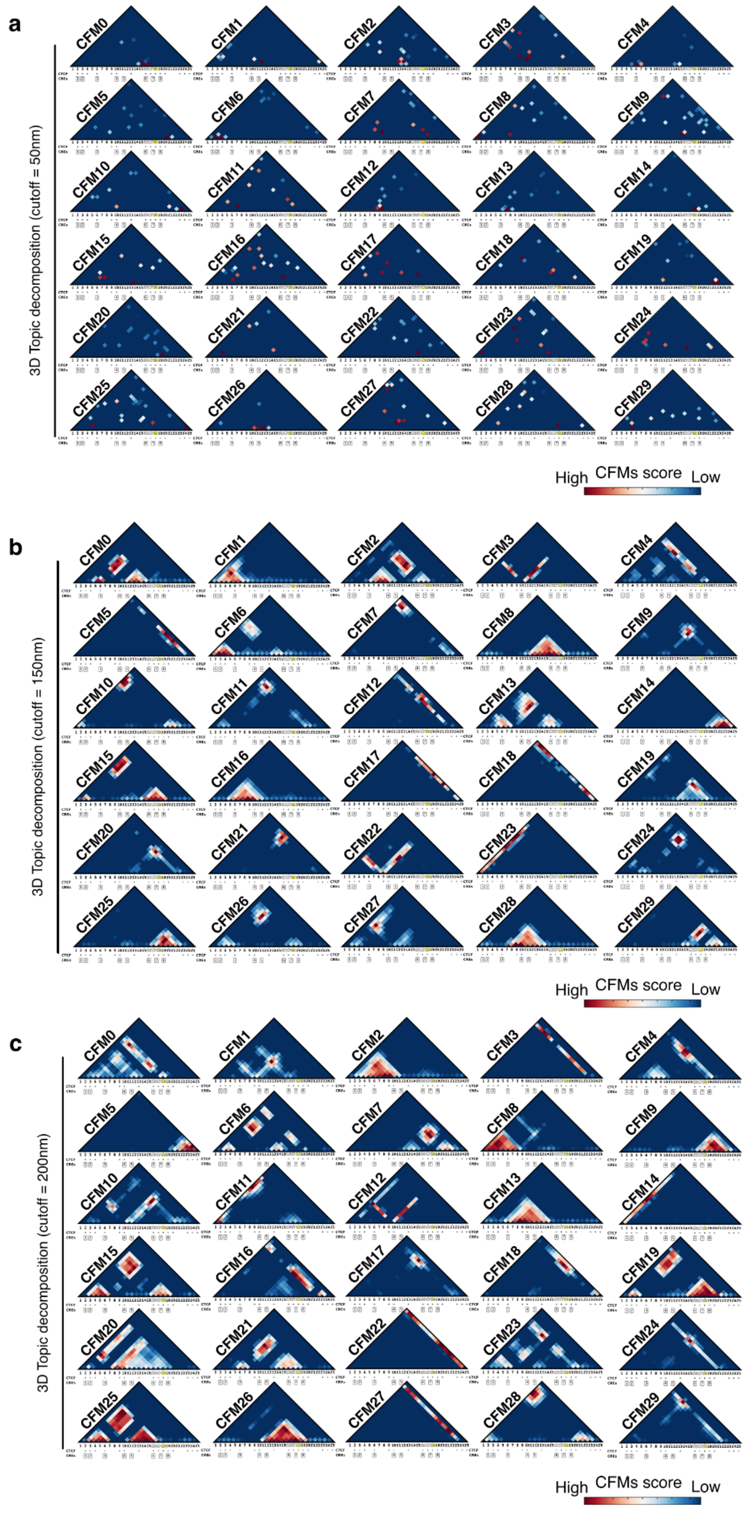


**a-c** 3Dtopic decompositions of *Pdx1* pancreas data using various cutoff distances: (**a)** 50 nm, (**b)** 150 nm, and (**c**) 200 nm.

### Supplementary Figure 4

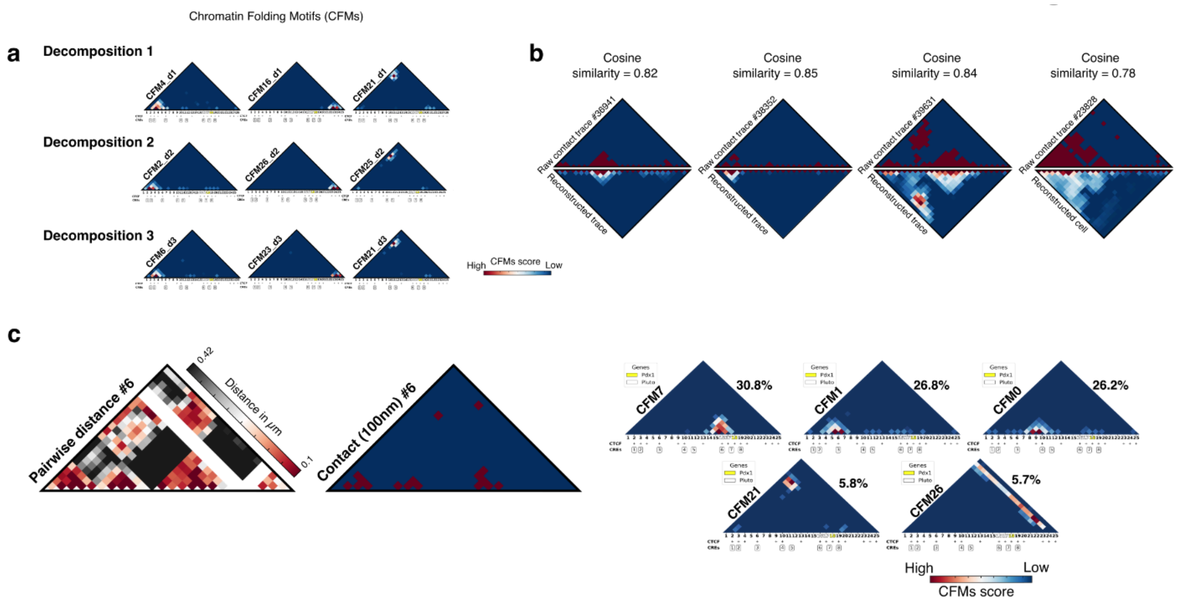


**a** Examples of three distinct CFMs identified in three independent decompositions.

**b** Side-by-side comparison of raw cell data and 3DTopic-reconstructed matrices on a model trained on pancreatic data along the Pdx1LR locus. Correlation between raw contact matrix (top) and reconstructed matrix (bottom) is calculated with Cosine similarity.

**c** Examples of raw cell data (pairwise distance matrix and contact matrix) and their most representative CFMs. Each CFM used to describe the corresponding cell is shown on the right, with its representativity score indicated alongside.

**Supplementary Figure 5**

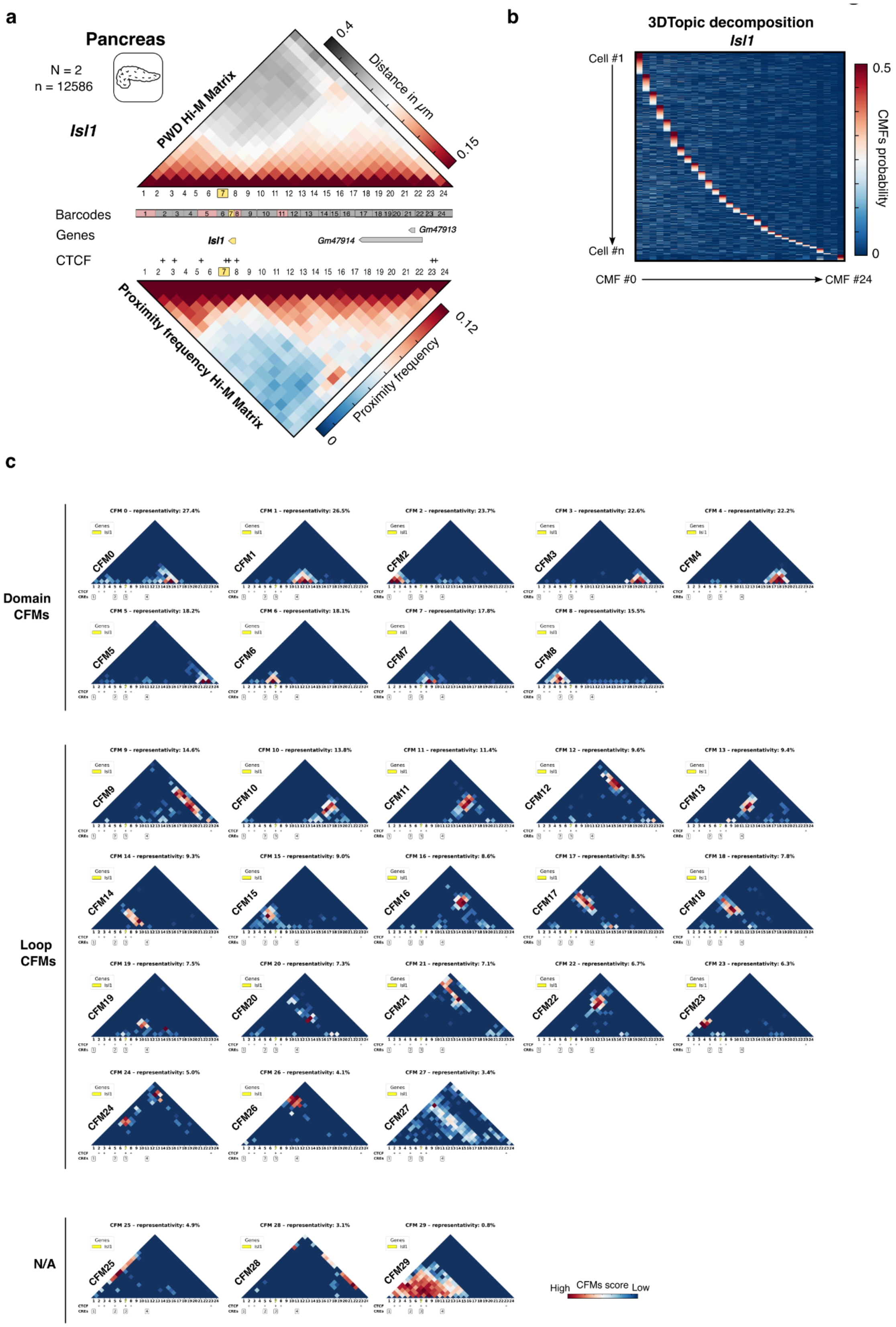


**a** Median Hi-M PWD matrix and Hi-M proximity map (100 nm cutoff distance) for the *Isl1* locus (chr13:116102031-116759941) in mouse pancreas constructed from 12586 traces from 2 experiments. Barcodes, genes, and CTCF sites are displayed along the locus.

**b** Probability distribution illustrating the decomposition of pancreatic cells into 30 CFMs along the *Isl1* locus.

**c** Complete gallery of chromatin folding motifs obtained from a decomposition along the *Isl1* locus classified in 2 distinct categories: Domains, Loops, with annotated CTCF and CREs sites.

### Supplementary Figure 6

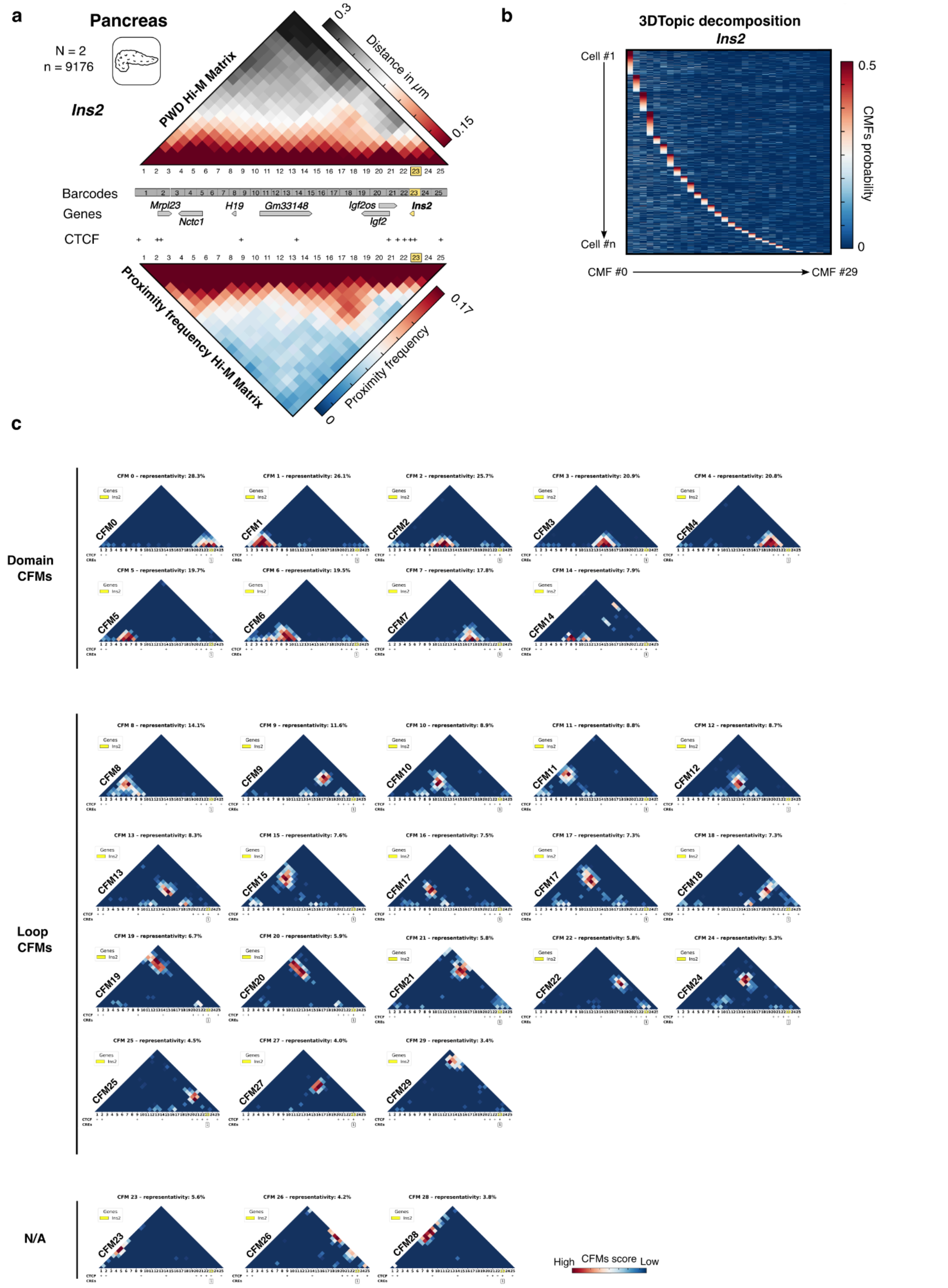


**a** Median Hi-M PWD matrix and Hi-M proximity map (100 nm cutoff distance) for the *Ins2* locus (chr7:142520024-142699856) in mouse pancreas constructed from 9176 traces from 2 experiments. Barcodes, genes, and CTCF sites are displayed along the locus.

**b** Probability distribution illustrating the decomposition of pancreatic cells into 30 CFMs along the *Ins2* locus.

**c** Complete gallery of chromatin folding motifs obtained from a decomposition along the *Ins2* locus classified in 2 distinct categories: Domains, Loops, with annotated CTCF and CREs sites.

### Supplementary Figure 7

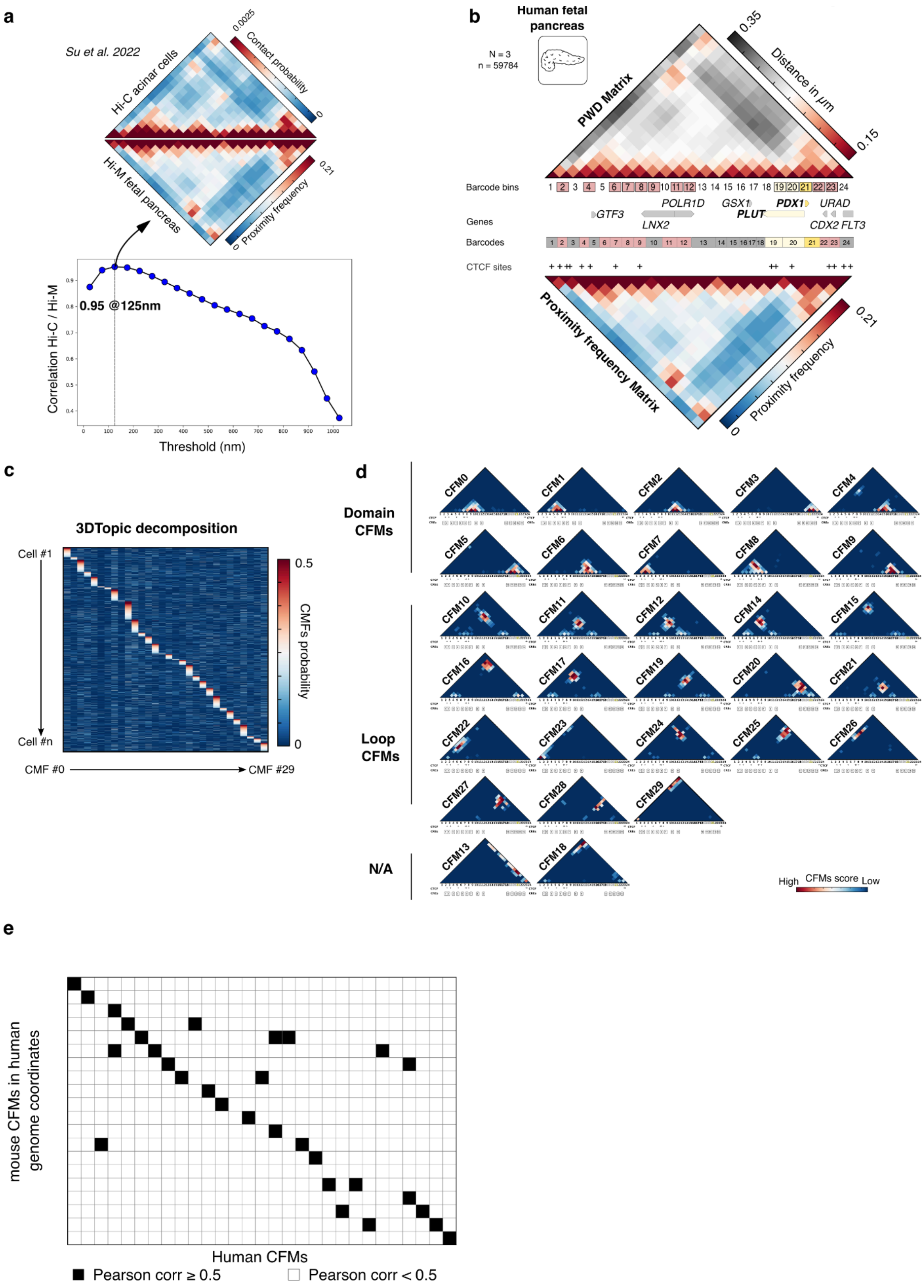


**a** Top: Comparison between the interpolated Hi-C contact matrix of human acinar cells for the *Pdx1* locus (chr13:27902436-28572561, hg19) from dataset GSE188311 (Su et al. 2022), and Hi-M proximity matrix from human fetal pancreas (cutoff distance = 125 nm). Bottom: Pearson correlation coefficient between the interpolated Hi-C contact map and Hi-M proximity frequency matrix for various cutoff distances. The highest correlation was achieved at a threshold of 125 nm, as indicated.

**b** Median Hi-M PWD matrix and Hi-M proximity map (125 nm cutoff distance) for the PDX1 locus (chr13:27902436-28572561, hg19) in human foetal pancreas constructed from 59784 traces from 3 experiments. Barcodes, genes, and CTCF sites are displayed along the locus.

**c** Probability distribution illustrating the decomposition of human pancreatic cells into 30 CFMs along the PDX1 locus.

**d** Complete gallery of CFMs obtained from PDX1 chromatin tracing data. CFMs are classified in two distinct categories: domains, loops, and N/A. Annotated CTCF and CREs sites are shown below.

**e** Matrix representing the binarized Pearson correlation between mouse CFMs mapped to the orthologous human genome coordinates (y-axis) and CFMs determined from experimental human PDX1 data (x-axis).

**Supplementary Figure 8**

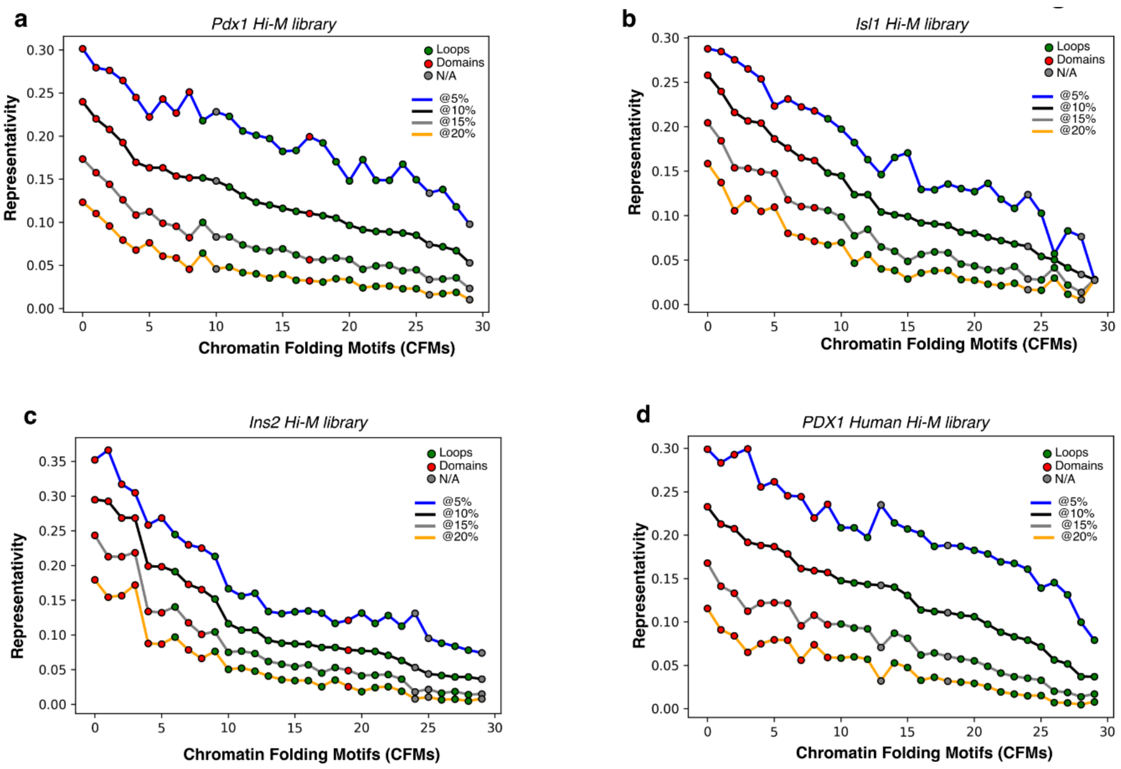


**a** Graph representing the global representativity of each CFM occurring in more than X% in pancreatic cells along the *Pdx1* locus, with X ranging from 5% (blue line), 10% (black line), 15% (gray line), to 20% (yellow line). Red circles represent domain CFMs, green circles represent loop CFMs, and gray circles represent CFMs that are not attributed to either loops or domains.

**b** Same as a, but for the *Isl1* Hi-M library.

**c** Same as a, but for the *Ins2* Hi-M library.

**d** Same as a, but for the human *Pdx1* locus.

### Supplementary Figure 9

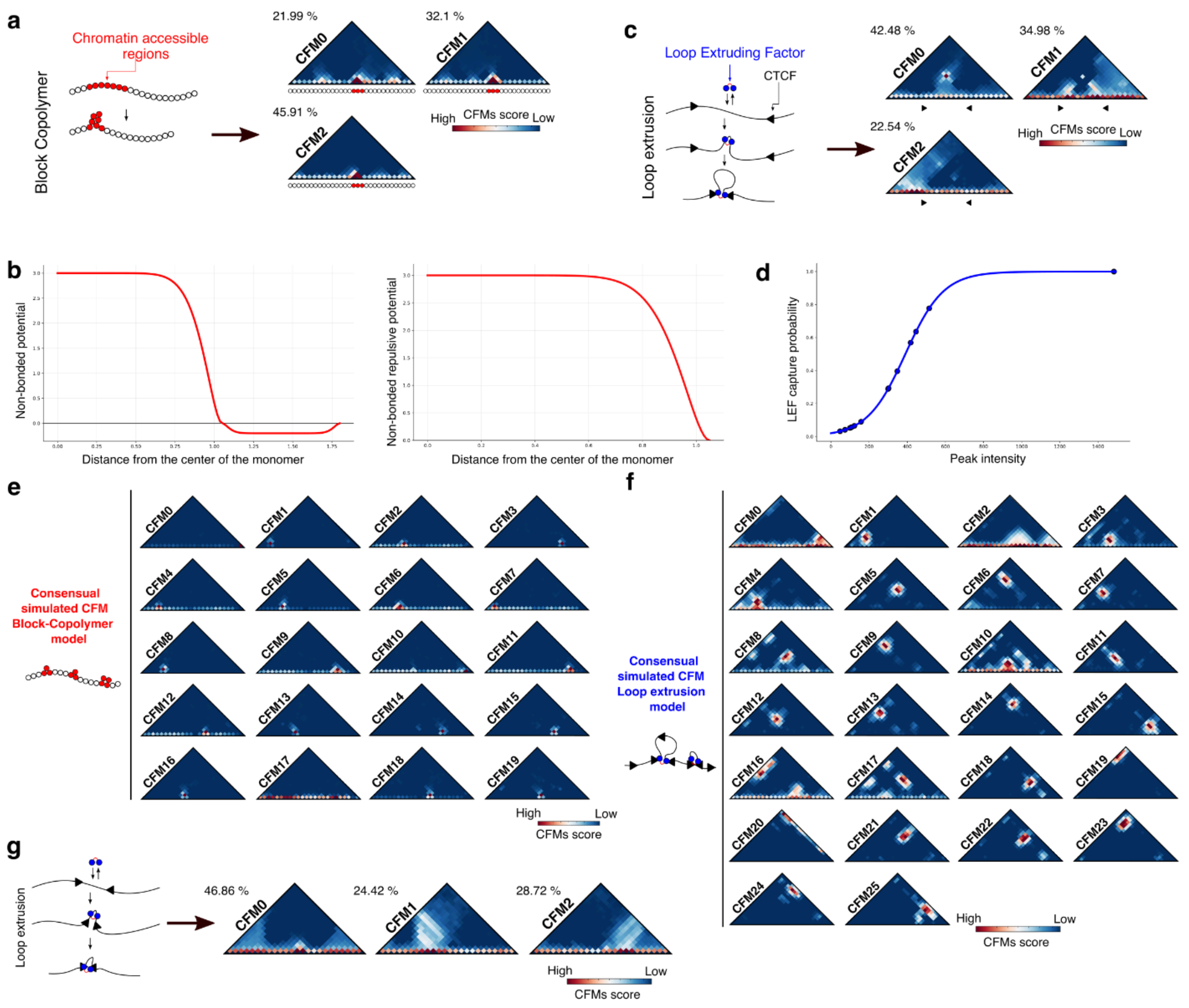


**a** (Left) Schematic diagram of block copolymer simulations where beads with ATAC-seq open peaks from pancreas (red) interact together when in close physical proximity (see Methods). (Right) Gallery of eight CFMs derived from the decomposition of simulated chromatin traces using the block copolymer model. Representativity for each CFM is shown on the top left.

**b** Potentials used to define forces between non-bonded monomers in the polymer chain during molecular dynamics simulations. Left: smoothed square-well potential for heteropolymers. Defines a common repulsive force between all types of monomers for excluded volume, and a soft attractive force only between monomers of the same type. Right: a truncated version of the above that only accounts for excluded volume in homopolymers. *Distances are given in bond length units.*

**c** (Left) Schematic diagram of a loop extrusion simulations (see *Methods*) of a chromatin region harboring CTCF sites (black triangles) assigned using experimental pancreas CTCF binding data. (Right) Gallery of eight CFMs derived from the decomposition of chromatin traces from loop extrusion simulations.

**d** Logistic curve used to compute LEF capture probabilities by CTCF based on the latter's peak intensity from CUT&Tag data. Curve mid-point $x_0$ = 390.231, steepness $k$ = 0.01.

**e** Complete gallery of consensual CFMs obtained from polymer simulation using a block copolymer model using the ATACseq peaks from pancreas at the Pdx1 locus from Liu *et al.*

(C. Liu et al., 2019) (see *Methods*).

**f** Same as c, but using a polymer model based on loop extrusion using the CTCF peaks from pancreas at the Pdx1 locus from Wang *et al.* (R. R. Wang et al., 2022) (see *Methods*).

**g** Loop extrusion simulations of two adjacent convergent CTCF sites located in adjacent bins (see *Methods*). 3DTopic reconstruction generates three CFMs: the major motif is a domain CFM, and the second and third correspond to alternative loop-extruded conformations. Percentages in the top left show the representativity of each CFM.

### Supplementary Figure 10

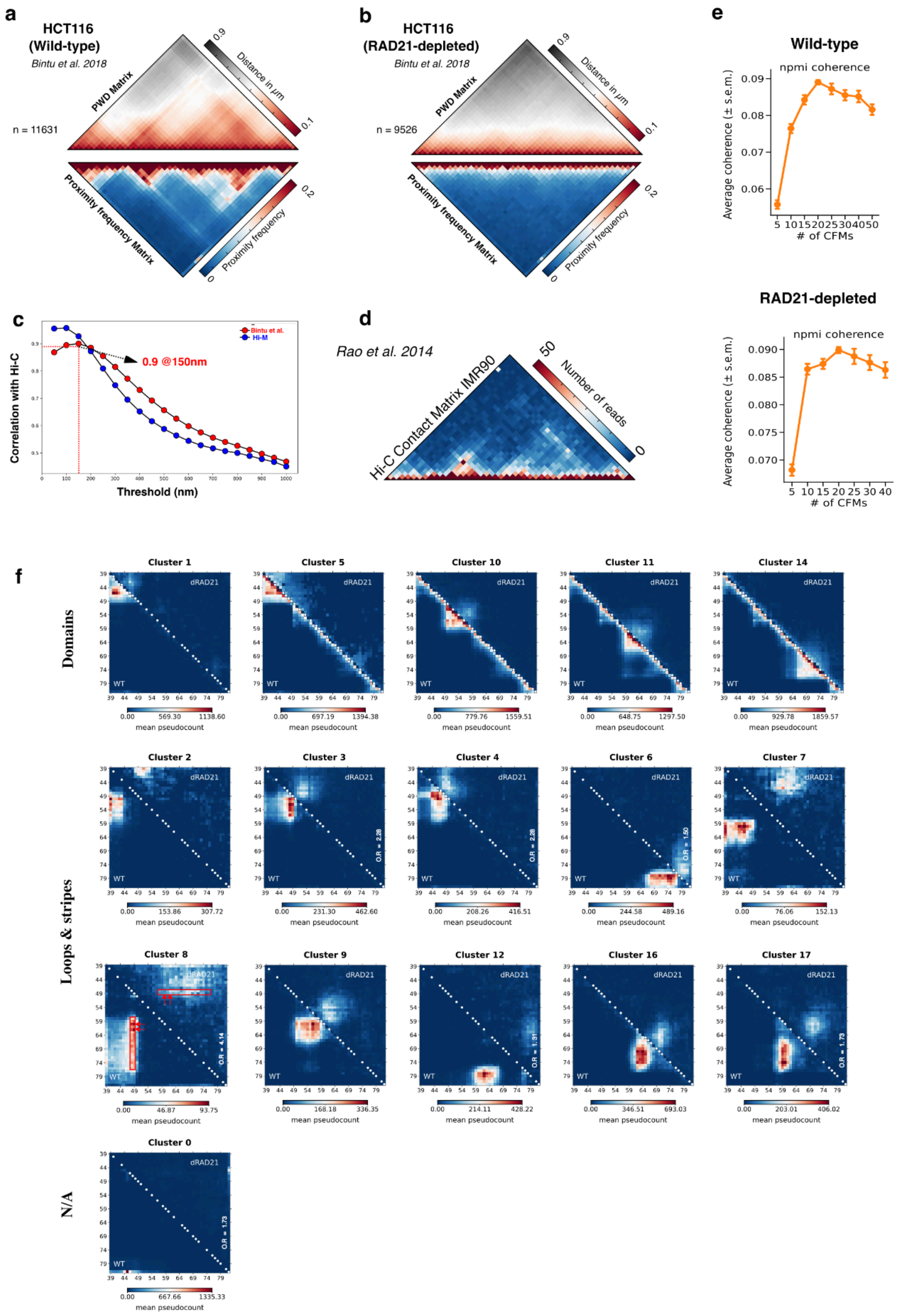


**a** Pearson correlation coefficients between the interpolated Hi-C matrix (GSE63525) and the proximity frequency matrices were computed across a range of distance thresholds for the (Bintu et al. 2018) dataset (red curve). These values are overlaid with the distribution of correlation values from our Hi-M data across different thresholds (blue curve). The highest correlation for this dataset was observed at a 150 nm threshold, while in our study, peak correlation occurred slightly below 100 nm.

**b** Hi-C contact matrix of IMR90 cells along the locus studied by Bintu *et al*, obtained from dataset GSE63525 (Rao et al. 2015).

**c-d** Median pairwise distance and proximity maps (150 nm cutoff distance) for (**c**) wild-type and (**d**) RAD21 depleted *HCT116* cells (*19*).

**e** Coherence scores from trainings performed for the wild-type and RAD21-depleted datasets as a function of the number of CFMs.

**f** Gallery of cluster-averaged CFM for each cluster in the wild-type UMAP (see **Fig. 3i**, left) showing comparisons between the wild-type (bottom left) and the RAD21-depleted (top right) datasets, white circles represent CTCF binding sites. Cluster-averaged CFMs are classified into domains, loops & stripes, or neither, based on visual inspection. In all cases, the characteristic domain and long-range interaction patterns observed in wild-type cells were either not present or markedly reduced after RAD21 degradation.

**Supplementary Figure 11**

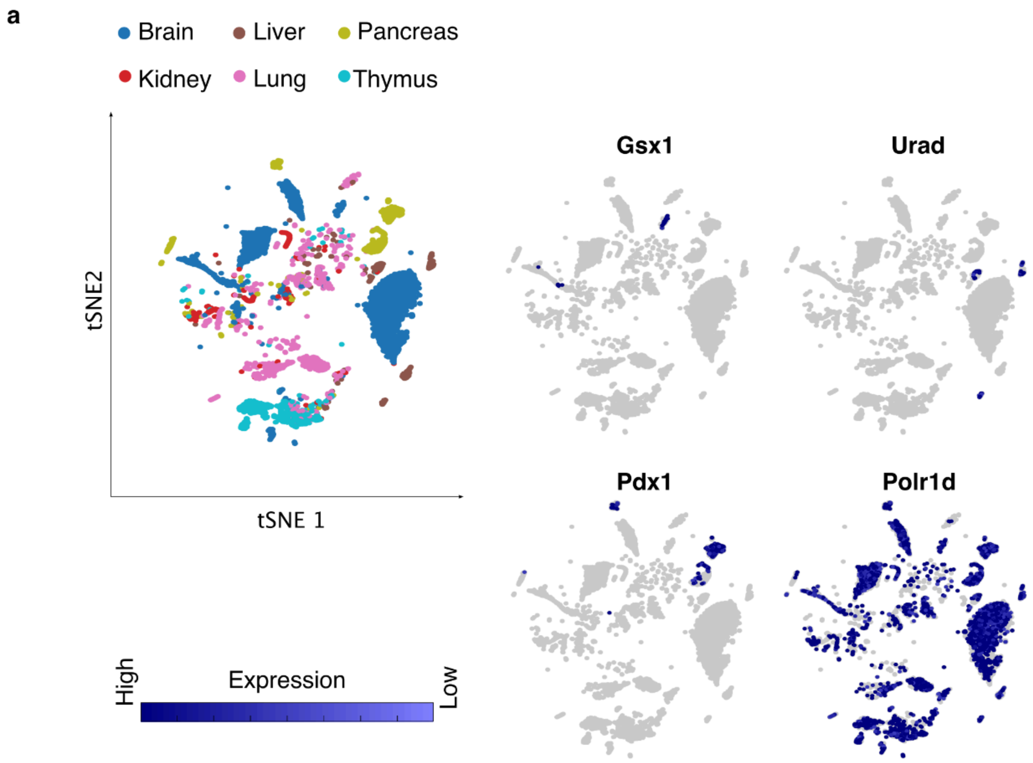


**a** Left: tSNE plot of cells from six different mouse tissues, color-coded by tissue type (see *Methods*). Right: tSNE plot displaying the expression levels of four different genes located along the *Pdx1* locus in single cells.

### Supplementary Figure 12

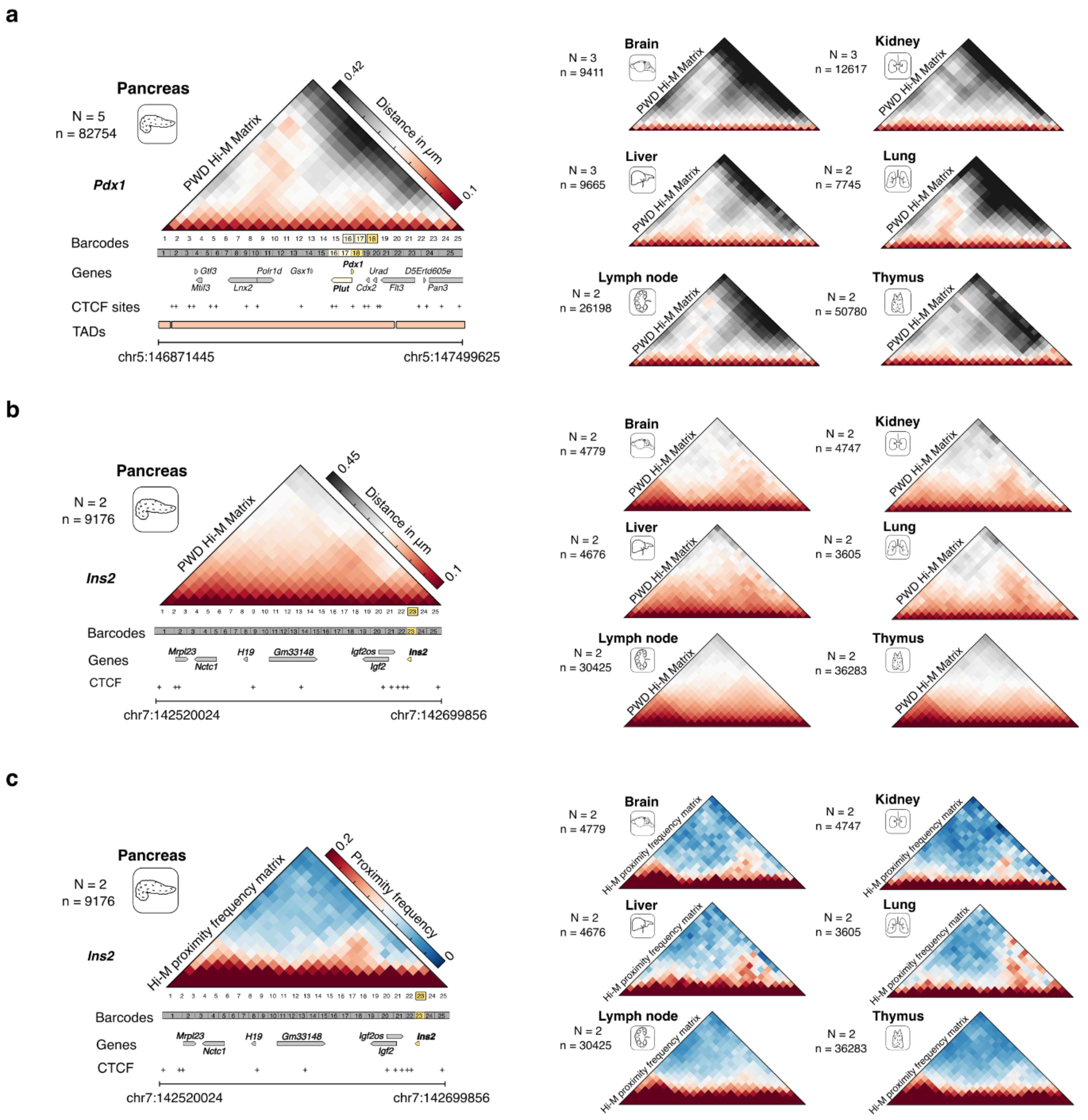


**a** Median PWD matrices for the *Pdx1* locus for Pancreas, Kidney, Lymph node, Lung, Liver, Thymus and Brain.

**b-c** Median PWD matrices (**b**) and Hi-M proximity (**c**) maps for the *Ins2* locus for the same tissues (cutoff distance: 100 nm).

**Supplementary Figure 13**

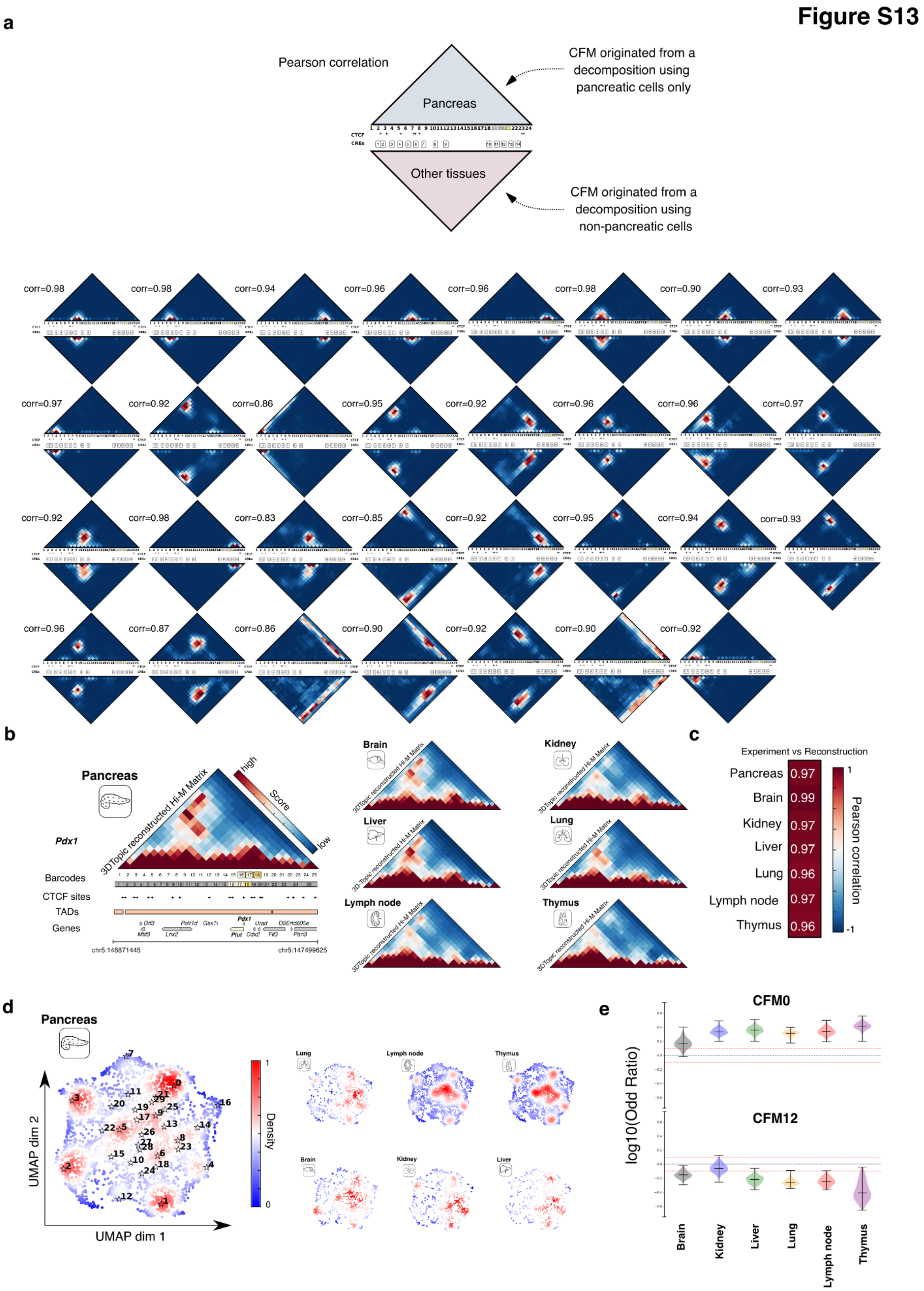


**a** Cluster-averaged CFMs were computed by averaging the CFMs for each cluster. Upper triangles represent cluster-averaged CFMs using decompositions from pancreas tracing data, while lower triangles show the cluster-averaged CFMs using decompositions using tracing data exclusively from all other tissues (brain, kidney, liver, lung, lymphnode, thymus). The Pearson correlations between the cluster-averaged CFMs from pancreas/other-tissues are shown above each map.

**b** 3DTopic reconstructed Hi-M matrix for the mouse pancreas and six other mouse tissues along the *Pdx1* locus.

**c** Matrix displaying the Pearson correlation between the experimental Hi-M contact matrix and the 3DTopic reconstructed Hi-M matrix for each of the seven tissues tested.

**d** Density distribution in the UMAP landscape for the pancreas and six other mouse tissues along the *Ins2* locus. Reference CFMs are displayed on the UMAP.

**e** Distribution of odds ratios from 500 independent decompositions for two CFMs, one domain CFM (0) and one loop CFM (12) using the pancreas as reference (see *Methods*). The dashed red line indicates log10(odds ratio) = 0.1 and -0.1, above and below which OR can be considered statistically significant.

### Supplementary Figure 14

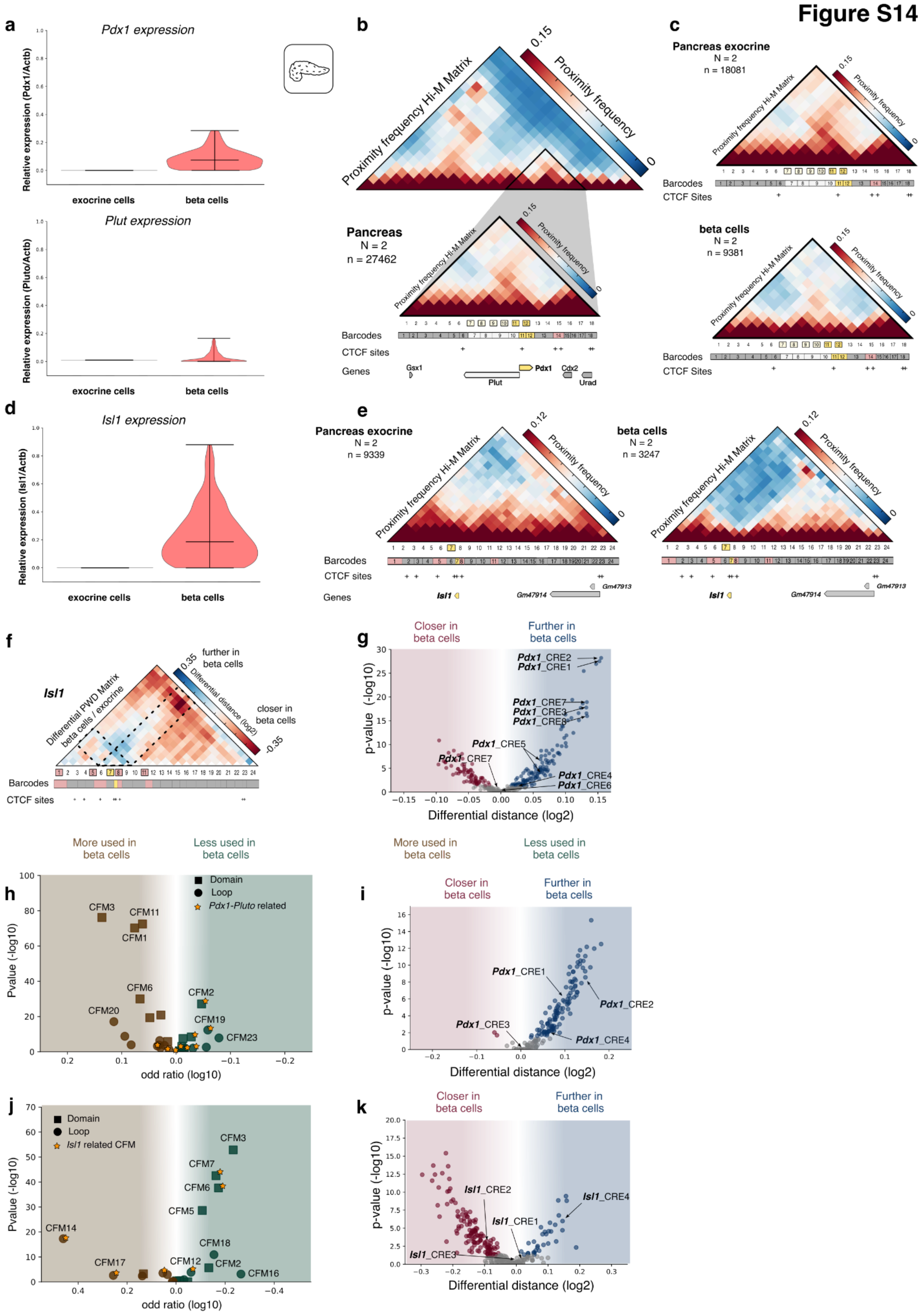

**a** Violin plot showing the relative expression of *Pdx1* or *Plut* compared to Actb in single cells in exocrine cells and in beta cells (see *Methods*), Top and bottom, respectively.

**b** Top: Hi-M proximity map along the *Pdx1* locus in mouse pancreas for the low resolution library. Solid triangle represents the genomic region imaged with the high-resolution library. Bottom: Hi-M proximity map along the *Pdx1* locus in mouse pancreas for the high resolution library (chr5:147180024-147324699, mm10) constructed from *n* = 27462 traces (*N* = 2 experiments, 2 different mice).

**c** Top: Hi-M proximity map along the *Pdx1* locus for the high resolution library in exocrine pancreas. Bottom: Hi-M proximity map in beta cells.

**d** Violin plot showing the relative expression of *Isl1* compared to Actb in single cells in exocrine cells and in beta cells (see *Methods*).

**e** Left: Hi-M proximity map along the *Isl1* locus in exocrine pancreas. Right: Median Hi-M proximity map in beta cells.

**f** Differential PWD Hi-M matrices between beta cells and exocrine cells for the Isl1 locus Hi-M library. Blue and red represent higher and lower distances in beta cells compared to exocrine cells, respectively. Dashed lines represent the bins occupied by Isl1t.

**g** Volcano plot showing the log2(beta cells/exocrine cells) PWD distance (x-axis) and -log10 (P-value) for the *Pdx1* locus. Gray circles indicate data points where the P-Value (y-axis) is below the 0.05.

**h** Volcano plot showing the log10 odd ratio (x-axis) between exocrine cells and beta cells, and -log10 (P-value) for each individual CFMs across 100 different decompositions. Domain CFMs are represented by squares, while loop CFMs are depicted by circles. Stars mark CFMs associated with *Pdx1* (Hi-M high-resolution library).

**i** Volcano plot showing the log2(beta cells/exocrine cells) PWD distance (x-axis) and -log10 (P-value) for the *Pdx1 HR* locus. Gray circles indicate data points where the P-Value (y-axis) is below the 0.05.

**j** Same as in h, for the *Isl1* locus.

**k** Same as in i, for the *Isl1* locus.

### Supplementary Figure 15

**Figure S15**

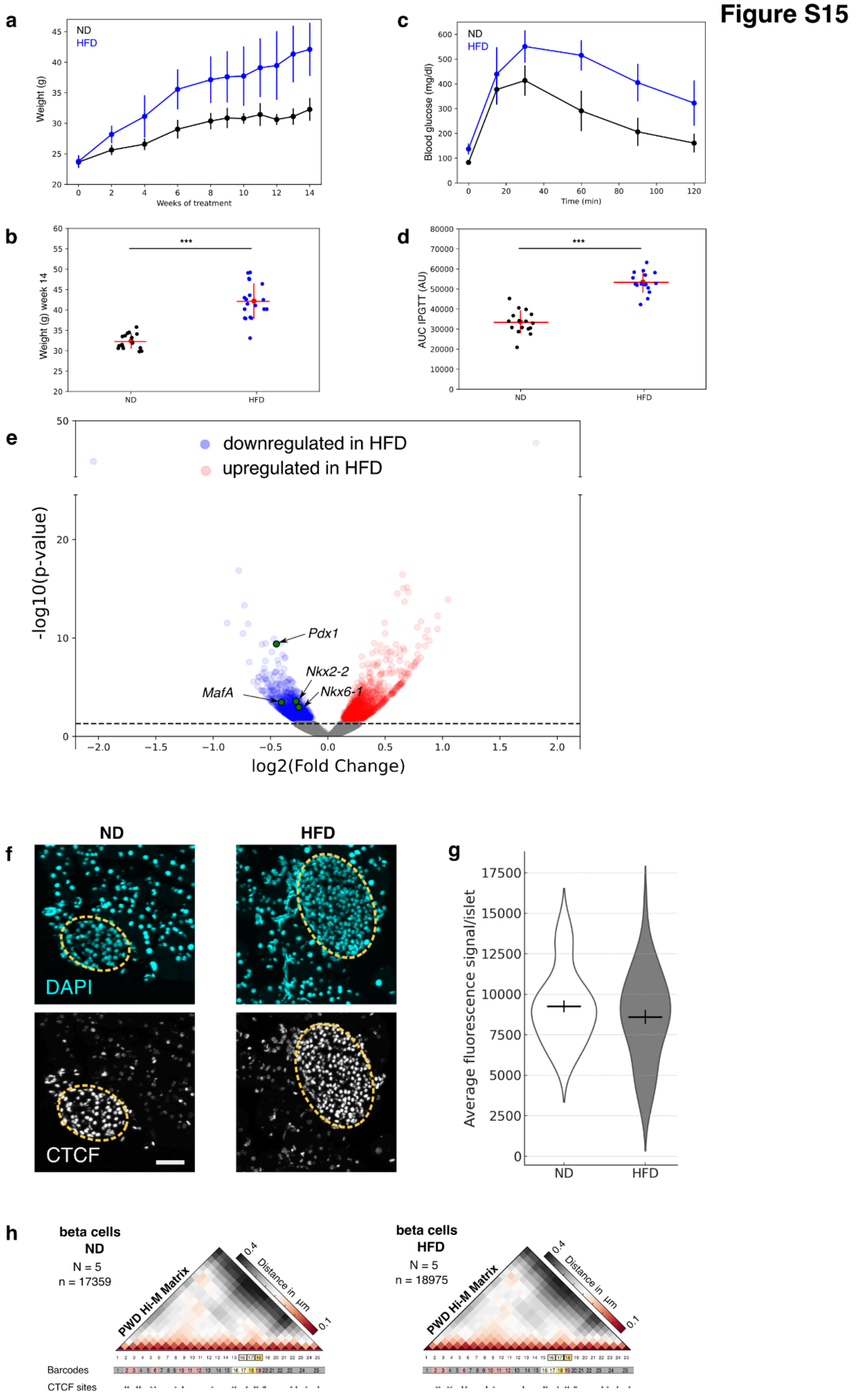

**a** Body weight (g) changes in mice fed with a normal diet (ND, black curve, n = 16) or a high-fat diet (HFD, blue curve, n = 19) over weeks.

**b** Body weights after 14 weeks of treatment for ND-fed and HFD-fed mice. P-value < 0.01, Mann-Whitney U test.

**c** Intraperitoneal glucose tolerance tests (IPGTT) in ND-fed and HFD-fed mice (see *Methods*).

**d** Area Under the Curve (AUC) analysis of IPGTT curve in c  P < 0.01, Mann-Whitney U test.

**e** Volcano plot showing genes that are differentially expressed (p-value < 0.05) in islets from high-fat diet (HFD)-fed mice compared to ND controls. Genes downregulated in HFD are shown in blue, and upregulated genes are shown in red. Transcription factors of interest : *Pdx1*, *MafA*, *Nkx2-2*, and *Nkx6-1*, which are significantly downregulated in HFD islets, are highlighted in green. Data from GSE131941 (Rosselot et al. 2019).

**f** Confocal images of pancreas frozen sections (10 µm) showing nuclei staining (DAPI in cyan, CTCF in white). ND: normal diet; HFD: high-fat diet. Islets are circled in yellow. Scale bar: 50 µm.

**g** Violin plot with mean and SEM of the average fluorescent signal in nuclei in islets after CTCF labeling (n = 49 (ND) and 55 (HFD) islets from N = 3 animals; Mann-Whitney U test, P = 0.42).

**h** Left: Median Hi-M PWD matrices along the *Pdx1* locus in beta cells under ND conditions. Right: under HFD conditions.

### Supplementary Figure 16

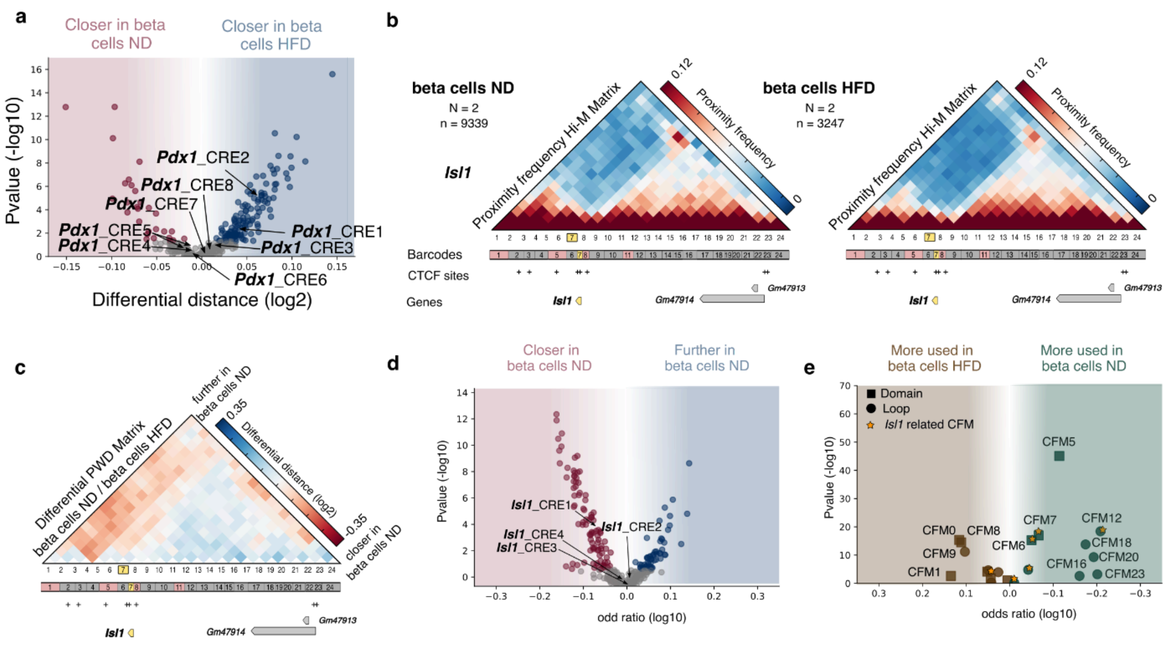


**a** Volcano plot showing the log2(beta cells ND/ beta cells HFD) PWD distance (x-axis) and -log10 (P-value). Gray circles indicate data points where the P-Value (y-axis) is below the 0.05.

**b** Left: Median Hi-M PWD matrices along the Isl1 locus in beta cells under ND conditions. Right: under HFD conditions.

**c** Differential PWD Hi-M matrices between beta cells ND and beta cells HFD. Blue and red represent higher and lower distances in beta cells ND compared to beta cells HFD, respectively.

**d** Volcano plot showing the log2(beta cells ND/ beta cells HFD) PWD distance (x-axis) and -log10 (P-value) for the *Isl1* Hi-M library. Gray circles indicate data points where the P-Value (y-axis) is below the 0.05.

**e** Volcano plot showing the log10 odd ratio (x-axis) between beta cells ND and beta cells HFD, and -log10 (P-value) for each individual CFMs accross 100 different decompositions. Domain CFMs are represented by squares, while loop CFMs are depicted by circles. Stars indicate CFMs related to the *Isl1* gene.
